# Structures of Nucleotide-Bound Redondovirus Rep Protein Link Conformation and Function

**DOI:** 10.1101/2025.08.25.672147

**Authors:** Saira Montermoso, Kushol Gupta, Ruth Anne Pumroy, Vera Moiseenkova-Bell, Frederic D. Bushman, Gregory D. Van Duyne

## Abstract

Circular Rep-encoding single-stranded DNA (CRESS-DNA) virus Rep proteins are multidomain enzymes that mediate viral DNA rolling-circle replication. Reps nick viral DNA to expose a 3’ end for polymerase extension, provide an NTP-dependent helicase activity for DNA unwinding, and join nicked ends to form circular viral genomes. Here, we present the first structures of a Rep protein from the *Redondoviridae* family, a newly discovered family of human-associated CRESS-DNA viruses that replicates within the oral protozoan *Entamoeba gingivalis*. Using cryo-EM, we characterized the hexameric structures of a Redondovirus Rep helicase bound with ATPγS, representing the initial ATP-bound state, and with ADP, reflecting the protein state after hydrolysis. The ADP state, but not the ATP state of Rep shows a staircase arrangement of DNA-binding loops that plays a central role in current models for SF3 helicase function. Additionally, we determined a head-to-tail dodecameric structure of ATPγS-bound Rep, in which both the helicase and endonuclease domains are ordered. Conservation of residues involved in stabilizing the dodecamer suggest that this assembly may be functionally relevant for many CRESS-DNA viruses. The positioning of endonuclease domains in the Rep hexamer, combined with our biophysical analyses of Rep oligomerization, provide new insights into Rep function during viral replication.

## Introduction

The circular Rep-encoding single-stranded DNA (CRESS-DNA) viruses are ubiquitous small viruses of Eukarya and Archaea (1). CRESS-DNA viruses have been isolated from a wide array of biomes, such as marine, plant, and human-associated communities (2). Many CRESS-DNA viruses appear to be benign commensals, while others are important pathogens, such as porcine circoviruses and geminiviruses (1). Redondoviruses are recently discovered CRESS-DNA viruses that were first identified in human respiratory samples (3, 4) and subsequently shown to be widespread in human oral and lung samples (5–7). Redondovirus recovery was increased in samples from human airway during acute illness, intubation and COVID19 infection (5, 8–12). Further studies showed that Redondovirus abundance was tightly correlated with that of the oral amoeba *Entamoeba gingivalis* (*13, 14*). A xenic culture of *E. gingivalis* was shown to harbor Redondoviruses, and Hi-C analysis showed that DNA of the two could be crosslinked, supporting *E. gingivalis* as the host of Redondoviruses (13).

According to present models, CRESS-DNA virus replication initiates following introduction of the viral genome into cells, where the single stranded viral DNA genome is copied into double stranded DNA by cellular enzymes (15, 16). The double stranded DNA is then the substrate for the viral-encoded Rep protein, which nicks the genome at extruded DNA cruciform structures formed at the origin of replication. Rep remains covalently joined to the DNA 5’-end through a phosphotyrosine linkage and the free 3’-end provides a substrate for a host cell DNA polymerase. The Rep protein is also an ATP-dependent DNA helicase, which is believed to assist in viral DNA replication via the rolling circle mechanism (17) and may also play a role in viral DNA packaging. Rep proteins are encoded by all CRESS viruses, and related Reps are encoded in some bacterial plasmids (1)

Despite the ubiquity and importance of CRESS-DNA viruses, understanding of CRESS Rep functional mechanisms is nascent. Rep has two functional domains (Fig. 1A). An amino-terminal HUH endonuclease domain is responsible for origin recognition, cleavage, and re-ligation to form ssDNA circles (18). The HUH domain is tightly coupled to an SF3 helicase domain that contains oligomerization and AAA+ ATPase subdomains (19). Structural models are available for related HUH endonuclease domains (20–22) and one structure has recently been described for a CRESS-DNA Rep helicase (23). The helicase model is a 3.8 Å cryo-EM structure of porcine circovirus type 2 Rep (pcRep) in an ADP-bound state. pcRep forms a hexamer with a narrow channel that can accommodate ssDNA, but not dsDNA. Two well-studied viral SF3 helicases are the large T antigen from Simian Virus 40 (SVTag) and the human papillomavirus E1 protein (24, 25). Compared to pcRep, the SVTag oligomerization subdomains form a larger channel that accommodates dsDNA, so that strand separation occurs between the domains or within the ATPase domains (26).

**Figure 1.**
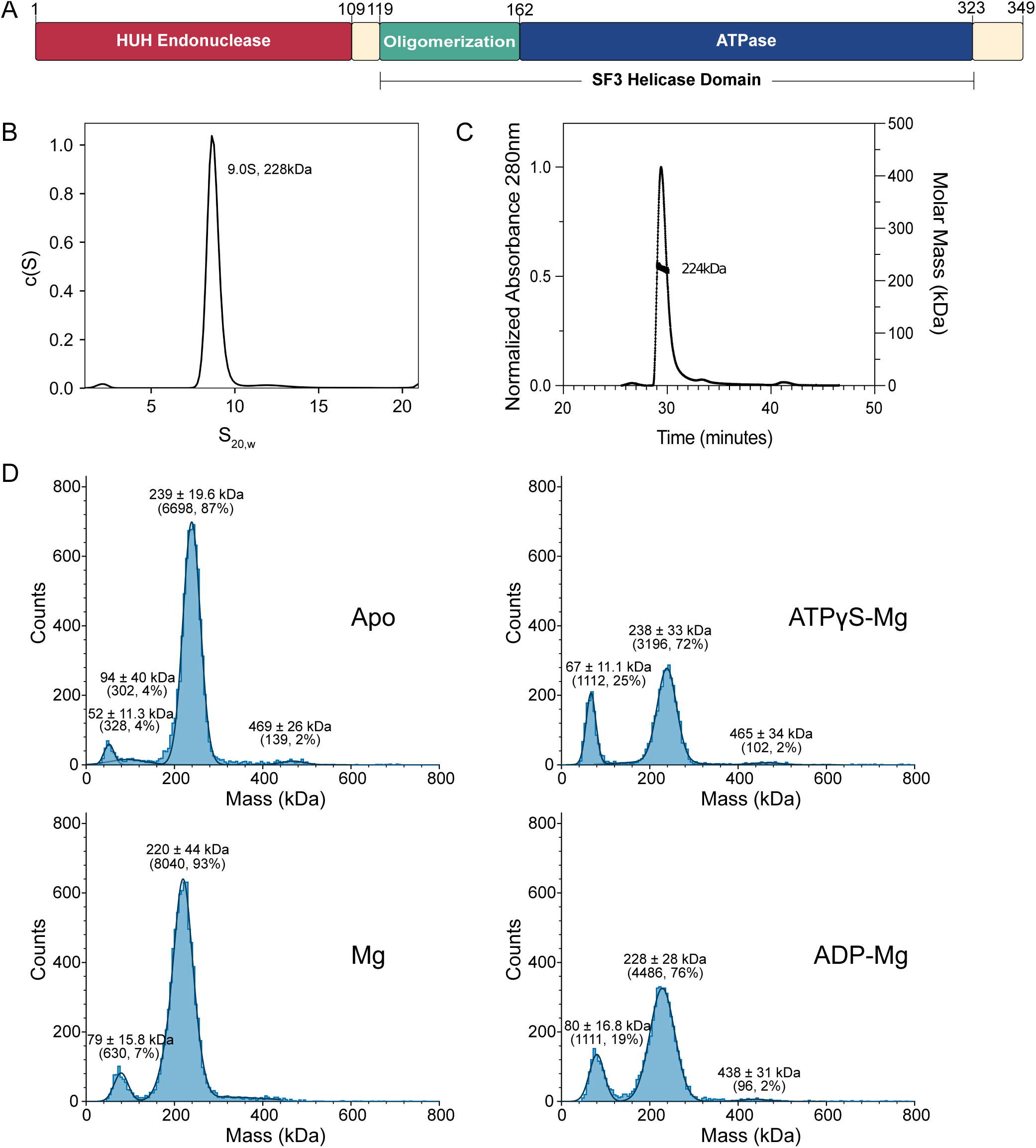
Properties of fbRep in solution. A. Domain architecture of fbRep. Rep consists of an N-terminal HUH-endonuclease domain (ED) and a C-terminal SF3 helicase domain. The helicase domain is comprised of oligomerization and AAA+ ATPase sub-domains. B. SV-AUC analysis of fbRep at 8.7 µM concentration. C. Size-exclusion chromatography in line with multi-angle light scattering (SEC-MALS) analysis of fbRep at 50 µM concentration. D. Mass photometry analyses of 100 nM apo-fbRep, and fbRep in buffers containing Mg^2+^, ATPγS + Mg^2+^, or ADP + Mg^2+^. Calculated masses of fbRep oligomers are monomer: 40.5 kDa; pentamer: 203 kDa; hexamer: 243 kDa; dodecamer: 486 kDa. All buffers contained 300 mM NaCl.

A recent cryo-EM analysis of SVTag reaction intermediates, combined with previous work in the same and related helicase systems, have established a mechanistic framework that is likely to be applicable to all hexameric helicases (25–27). In this “symmetric rotary mechanism”, DNA-binding β-turns of the ATPase subunits form a “spiral staircase” that promotes unidirectional movement of ssDNA through the central channel, and the complementary strand is extruded from the hexameric assembly. The staircase is dynamically manipulated by hydrolysis of ATP at the subunit interfaces. Indeed, the pcRep cryo-EM structure has a staircase arrangement of DNA-binding loops in support of this model (23). Although the mechanism of ssDNA translocation by the conserved ATPase domains is expected to be similar among SF3 helicases, the functions of CRESS-DNA Rep proteins differ in many ways from the origin opening roles of SVTag and E1 (28, 29), and the CRESS Rep functions remain incompletely understood.

Here we present a structural study of the Rep protein of a Vientovirus, a viral species in the *Redondoviridae* family (30). Previously Vientovirus Rep protein was purified and shown to be capable of site-specific cleavage at a DNA stem-loop designed to mimic the inferred origin of viral replication (7). We report that Rep protein from Vientovirus FB (fbRep, Fig. 1A) is a tractable model for understanding CRESS-DNA virus Rep mechanisms at the structural level. We first established the oligomeric state of fbRep using biophysical methods, identifying pentamers, hexamers and dodecamers across a broad range of concentrations and ionic strengths. We then generated cryo-EM structures for hexameric fbRep bound with ATPγS at 2.36 Å and bound with ADP at 3.20 Å, revealing the allosteric changes in the ATPase sub-domains that link the two nucleotide bound states. We also report the structure of a novel dodecameric form of fbRep•ATPγS composed of head-to-tail hexamers where the HUH endonuclease domains of one hexamer are fully resolved. These models provide important new insights into Rep function and will facilitate the design of experiments to test Rep functions in CRESS-DNA virus biology.

## Results

### Oligomeric properties of Redondovirus Rep

In prior work, we established that recombinant full-length fbRep was capable of carrying out DNA nicking and joining activities in vitro (7). However, those protein preparations were purified and stored in 1 M NaCl, which is poorly suited for structural studies. In addition, we found that fbRep exists primarily as pentamers in high salt by analytical ultracentrifugation (AUC) (Supplemental Figure 1). We therefore re-optimized our purification protocol and assessed the monodispersity of fbRep at salt concentrations closer to physiological levels using sedimentation velocity-AUC (SV-AUC) (Fig. 1B). In 300 mM NaCl and at 8.7 µM concentration, we observed a single 9.0 S species with a mass of ∼228 kDa (calculated monomer 40.5 kDa, pentamer 203 kDa, hexamer 243 kDa). Using size-exclusion chromatography in-line with multi-angle light scattering (SEC-MALS), the absolute mass determined was 224 kDa ± 5%, consistent with a mixture of pentamer and hexamer forms (Fig. 1C).

We extended our analyses to include fbRep in the presence of ADP and ATP (or ATPγS) across a broad range of Rep concentrations. We first applied mass photometry to assess the impact of nucleotides on the oligomeric state of fbRep at physiological concentrations (Fig. 1D). Hexamers are the predominant species at 100 nM concentration in the apo, Mg, ADP-Mg, and ATPγS-Mg forms, with evidence for dissociation into monomers and/or dimers that are more prevalent in the presence of nucleotides. To further probe the solution properties of nucleotide-bound fbRep, we employed size-exclusion chromatography in-line with synchrotron small-angle X-ray scattering (SEC-SAXS) (Supplemental Figures 2 & 3). The scattering profiles were analyzed using singular value decomposition with evolving factor analysis (SVD-EFA), which identifies the scattering contributions from the major species present in solution (31). For apo-fbRep and fbRep•ADP at ∼100 µM concentration, we found evidence for both Rep hexamers and a larger species whose mass corresponds to a dimer of Rep hexamers (Table 1). We also found that apo-Rep vs. Rep•ADP hexamers show distinct P(r) profiles, indicating changes in quaternary structure when nucleotide binds (Supplemental Figure 2). For fbRep•ATP at the same concentration, we observed only the larger, dodecameric species (Table 1).

**Table 1.** Properties of Rep derived using SEC-SAXS with SVD-EFA analyses.

| Nucleotide State | Guinier |  | GNOM |  | Mass (kDa) | Oligomer |
| --- | --- | --- | --- | --- | --- | --- |
| | $qR_g$ | $R_g$ (Å) | $R_g$ (Å) | $D_{max}$ (Å) | By $V_c$ | |
| apo-fbRep | 0.42 – 1.29 | $52.4 \pm 0.3$ | $52.3 \pm 0.1$ | 155 | 553 | Dodecamer |
| | 0.48 – 1.39 | $47.8 \pm 0.6$ | $47.2 \pm 0.6$ | 139 | 286 | Hexamer |
| +1 mM ADP | 0.42 – 1.29 | $54.8 \pm 0.4$ | $53.1 \pm 0.1$ | 239 | 476 | Dodecamer |
| | 0.40 – 1.28 | $49.2 \pm 0.5$ | $49.1 \pm 0.2$ | 162 | 263 | Hexamer |
| +1 mM ATP | 0.48 – 1.41 | $54.4 \pm 0.3$ | $54.1 \pm 0.2$ | 177 | 510 | Dodecamer |

### Cryo-EM structure of Redondovirus Rep bound by ATPγS

To obtain the structure of fbRep in the ATP-bound state, we prepared Rep complexes using the non-hydrolyzable analog ATPγS. Preliminary experiments indicated that fbRep was prone to aggregation at salt concentrations less than 300 mM. In 300 mM NaCl, cryo-EM micrographs displayed fbRep oligomers as well-defined and easily recognizable particles. Single particle analysis of fbRep•ATPγS revealed a diverse composition of pentamers, hexamers, and double hexamers (Supplemental Figure 4). Hexamers were the most prevalent form observed (examples in Fig. 2A), followed by pentamers.

**Figure 2.**
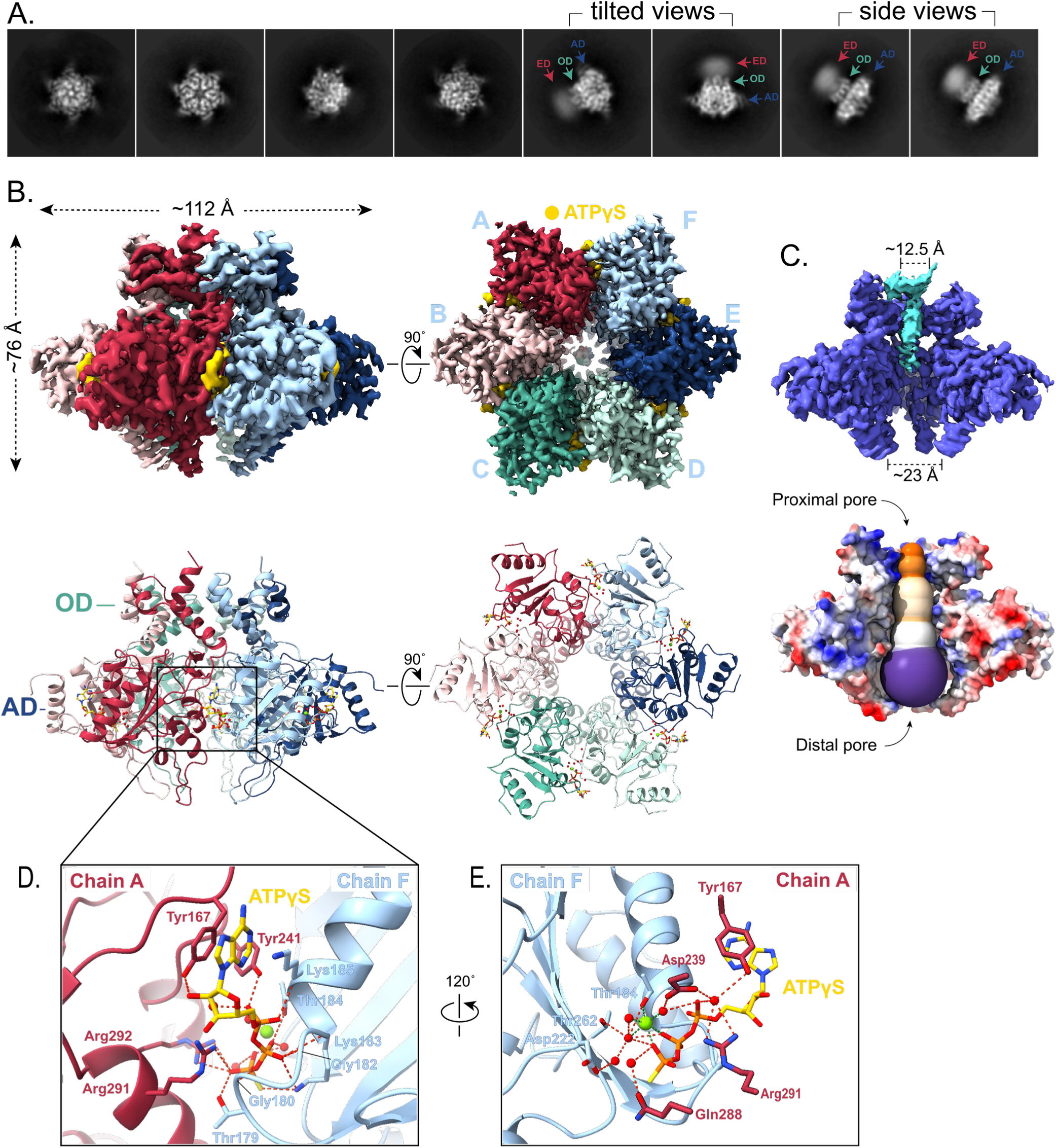
Cryo-EM structure of ATPγS-bound fbRep hexamer. A. Representative 2D class averages of fbRep hexamers. Diffuse density for the endonuclease domains (ED) is indicated in side and tilted views. B. Orthogonal views of reconstructions for the ATPγS-bound fbRep hexamer are shown on top with corresponding cartoon representations below. Bound ATPγS is drawn in yellow. Oligomerization (OD) and ATPase (AD) subdomains of Rep are indicated. The endonuclease domains could not be fit. C. Views of the fbRep•ATPγS helicase pore channel. Blue density in a cutaway of the reconstruction may represent nucleic acid captured during expression and purification. A solvent-accessible profile of the channel reveals a uniform pore with a relatively narrow proximal pore through the oligomerization domains that gradually widens to the distal pore at the bottoms of the ATPase domains. D. The ATP binding site is located between ATPase domain subunits. E. Rotated view of the ATP binding site showing coordination of magnesium ion and surrounding water molecules.

Three-dimensional classification of the hexameric particles led to the reconstruction of two distinct fbRep hexamers (Supplemental Figure 5, Table 2). The first was reconstructed from two similar hexamers composed of 661,708 particles. High-resolution 3D refinement, without internal symmetry imposed, resulted in a 2.36 Å reconstruction of a hexamer that features highly resolved helicase domains with no visible endonuclease domain density (Fig. 2A). The second hexamer reconstruction featured visible, but diffuse endonuclease domain density. This map, however, refined to lower resolution (3.19 Å) with an endonuclease domain density that could not be interpreted.

**Table 2.**
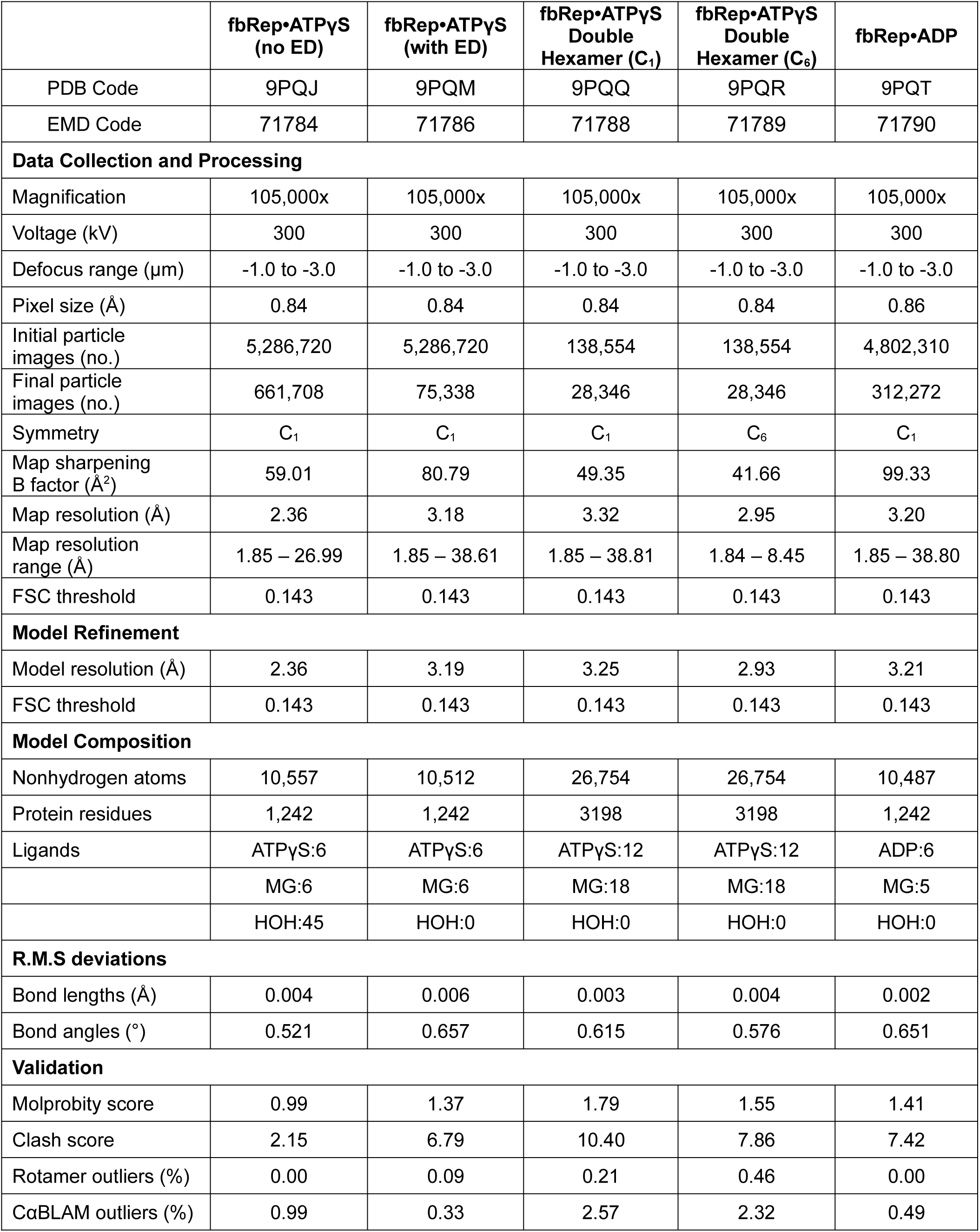
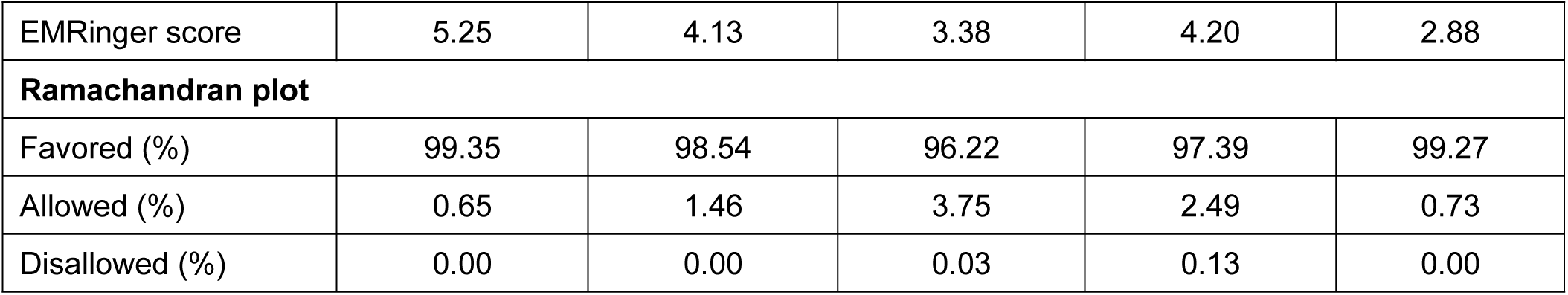
Cryo-EM data collection and model statistics.

The 2.36 Å cryo-EM map allowed for the facile building and refinement of an atomic model revealing a six-fold symmetric ring assembly of oligomerization and ATPase sub-domains with overall dimensions of ∼72 x 112 Å (Fig. 2B). The six nucleotide binding sites located between adjacent ATPase subunits are fully occupied and could be fit with high fidelity. The map also revealed weak density in the central pore formed by the oligomerization subdomains (Fig. 2C). A similar density was observed in the pcRep•ADP cryo-EM map (23), but in that case the additional density extended well into the ATPase subdomains and could be modeled as heterogeneous ssDNA that was carried over from *E. coli* expression and purification. The density in fbRep•ATPγS is also likely to be *E. coli*-derived nucleic acid but is present at a lower occupancy and we did not attempt to model it. The pore formed by the oligomerization subdomains is ∼12 Å in diameter and remains constricted into the proximal pore formed by the ATPase subdomains. The channel opens at the distal end of the ATPase hexamer, with a diameter of ∼23 Å (Fig. 2C).

The high resolution fbRep•ATPγS structure provides a detailed model for the ATP-specific interface formed between adjacent helicase subunits. These interactions, along with alternative interactions in the corresponding ADP-specific interface, control the ATPase domain movements and consequent positioning of the DNA-binding loops responsible for staircase formation and helicase activity (25, 26). A close-up view of the ATP interface between subunits A and B is shown in Fig. 2D; the other five interfaces are nearly identical. The triphosphate of ATPγS is buried in the inter-subunit interface, along with a magnesium ion and several tightly bound water molecules (Fig. 2E). The adenosine fragment is nestled between two loops and an α-helix of adjacent subunits and is partly solvent exposed. ATP binding to fbRep resembles a key inserted into a keyhole, with the triphosphate ‘key’ matching the internal lock mechanism of the ATPase domains. In addition to interactions involving the canonical Walker A motif (Thr179, Gly180, Gly182, Lys183, and Thr184) and the arginine finger (Arg291, Arg292), the Lys185 amide, the Tyr241 hydroxyl, and the Tyr167 hydroxyl also make hydrogen bonded interactions to ATPγS. The buried Mg^2+^ ion is coordinated by the β- and γ-phosphates of ATPγS, Thr184 hydroxyl, as well as three water molecules. Details of fbRep interactions with ATPγS and Mg^2+^ are provided in Supplemental Figure 6.

The DNA-binding β-turns of the fbRep•ATPγS structure do not form a spiral staircase arrangement; they are approximately co-planar and share the C_6_ symmetry of the oligomerization subdomain. A similar arrangement was observed in the SVTag•ATP hexamer structure (27) and is likely due to the rigidity of the ATP interfaces combined with the lack of a ssDNA substrate to provide a staircase interaction template. To quantitate this structural feature and provide a means of comparison to other structures, we transformed the fbRep hexamer coordinates to an inertial frame with Cartesian Z coincident with the symmetry axis defined by the oligomerization subdomains. The z-coordinates of Gly249 in the β-turn ‘step’ then provide the helical pitch, if any, of the staircase (Gly249 in fbRep is the structural homolog of Gly240 in pcRep and His513 in SVTag). This analysis for fbRep•ATPγS confirms that the DNA-binding loops are co-planar (Table 3).

**Table 3.**
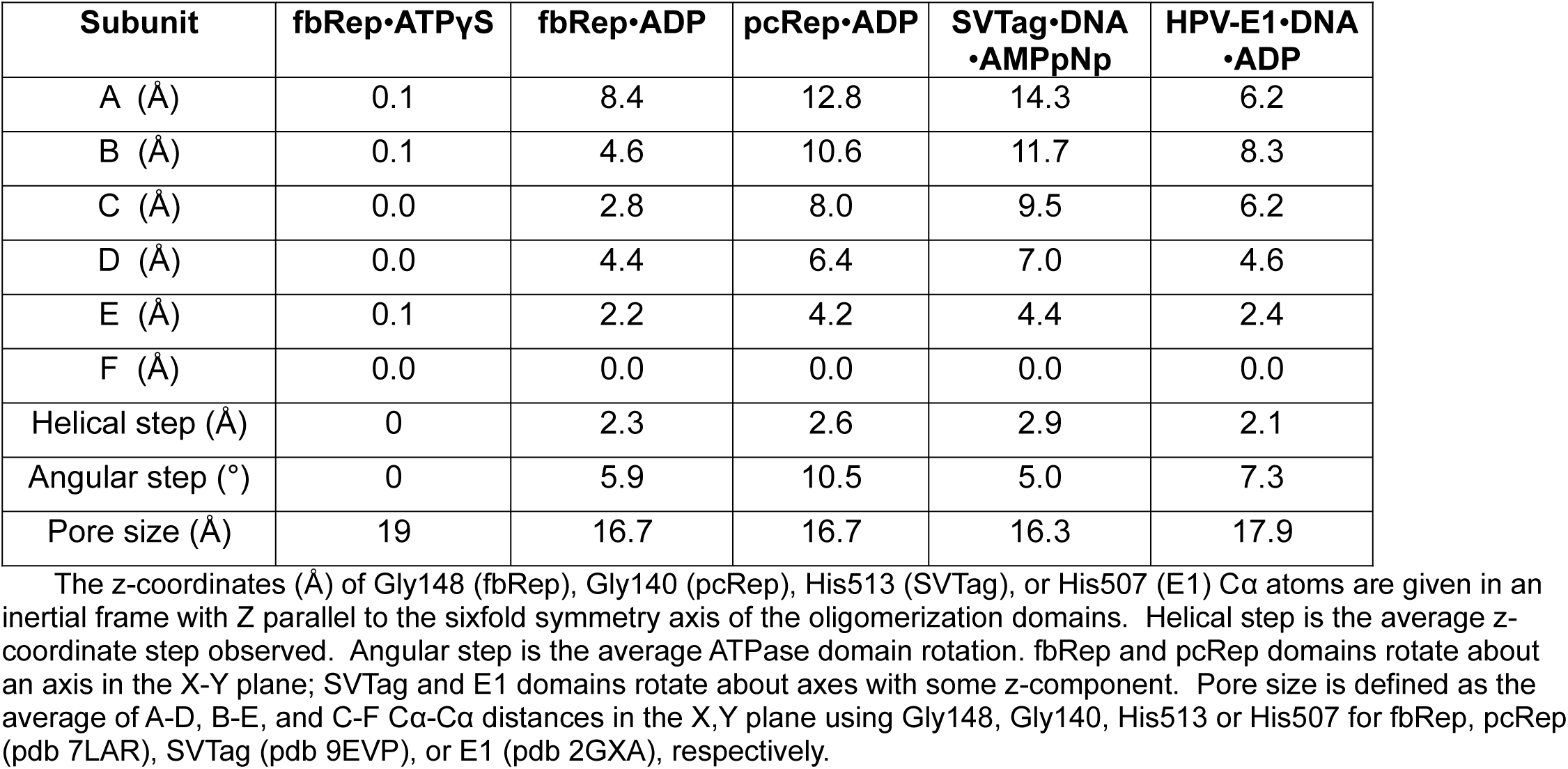
Helical parameters for fbRep and other SF3 family helicases.

### Cryo-EM structure of Redondovirus Rep bound by ADP

We prepared ADP-bound fbRep in a similar manner to ATPγS-bound fbRep and collected images for single particle analysis. In the ADP-bound form, hexamers were also the most abundant Rep oligomers observed in 2D class averages (Fig. 3A). The fbRep•ADP structure was determined using 312,272 particles with an overall resolution of 3.20 Å (Table 2, Supplemental Fig. 7). Overall, the ADP-bound hexamer resembles the ATPγS form (Fig. 3B, Supplemental Fig. 8), but closer inspection reveals that the similarity is primarily confined to the oligomerization domains, which are nearly superimposable with an r.m.s. difference of 0.83 Å. The quaternary structure of the ATPase domains is different in the ADP-bound structure, with an increased distal pore diameter of 45 Å the most conspicuous feature (Fig. 3C).

**Figure 3.**
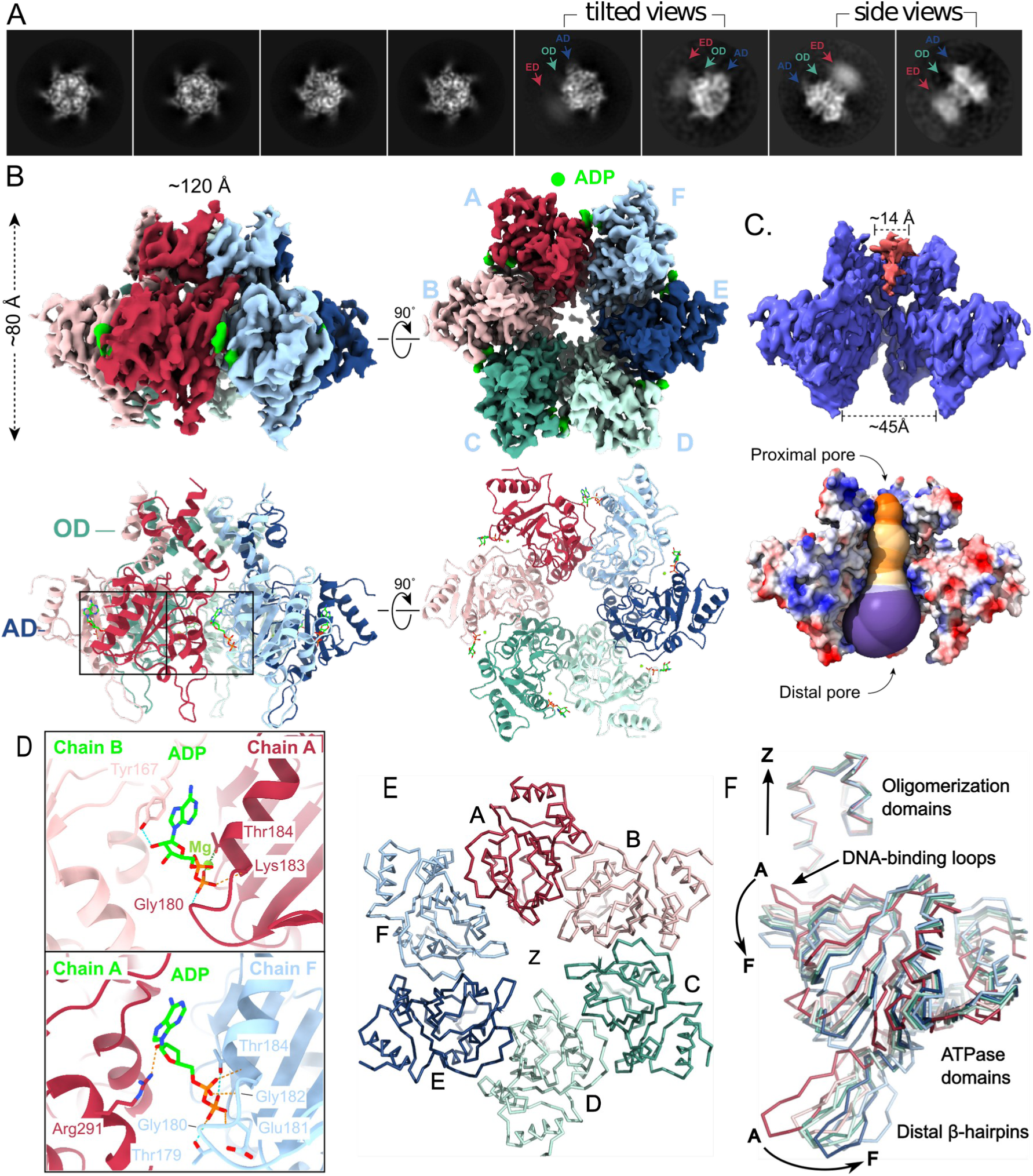
Cryo-EM structure of ADP-bound fbRep hexamer. A. Representative 2D class averages with diffuse density for the endonuclease domains indicated. B. Orthogonal views of the ADP-bound fbRep hexamer reconstruction with ADP density shown in lime green. Corresponding ribbon representations are shown below. The endonuclease domains could not be fit. C. Views of the pore channel. Pink density in the oligomerization domain channel is likely nucleic acid captured during expression and purification. The solvent-accessible profile of the channel generated using MOLEonline reveals a varying pore channel different than the pore channel profile of fbRep•ATPγS. The variations observed can be attributed to varying degree of ATPase domain movements of the helicase subunits. D. Examples of two of the six unique ADP binding sites. Although the diphosphate-Rep interactions are maintained in each interface, the remaining interactions differ with fewer total interactions compared to ATPγS. E. Spiral staircase formed by the DNA-binding loops of the ATPase domains. The oligomerization domains have been cut away for clarity. View is down the symmetry (Z) axis. The loops form a right-handed downward spiral going from subunit A to F (see also Table 3). F. Superposition of the six oligomerization domains. A successive rotation of ATPase domains A-F results in movement of the DNA-binding loops downwards and movement of the distal β-hairpins outwards. The sixfold symmetry axis is aligned with Cartesian Z as indicated.

ADP ligands could be readily fit into the six nucleotide binding site densities. ADP binds primarily to one subunit, where the same interactions are formed in each case. There are fewer interactions formed with neighboring subunits compared to ATPγS and those interactions vary among the six interfaces (Fig. 3D, Supplemental Fig. 9). The ADP interface contributes weaker constraints on the quaternary structure of the hexamer, allowing the ATPase domains to undergo small, successive rotations from subunit A to subunit F. These rotations generate a helical arrangement of DNA-binding loops around the pseudo-symmetry axis of the hexamer (Figs 3E & 3F). A z-coordinate analysis of the DNA-binding loops shows a clear spiral staircase arrangement, with the loops spanning 8 Å in the Z direction and an average helical step size of 2.3 Å (Table 3). The ATPase subunits rotate about an axis in the X-Y plane that passes close to the Cα atom of Tyr199, moving the DNA-binding loops away from the oligomerization domain pore (i.e., lowering the z coordinate) and expanding the distal pore formed by the ATPase domains (Fig. 3F). The staircase arrangement of DNA-binding loops decreases the proximal pore size by about 2 Å, going from the symmetric ATPγS structure to the asymmetric ADP form (Table 3).

A similar staircase configuration was observed in the pcRep•ADP structure (23), where we have calculated a 12 Å z-range, a 2.6 Å average helical step, and an identical pore size (Table 3). Unlike the pcRep structure, however, we do not observe strong density within the ATPase subunit channel that could be interpreted as ssDNA. Thus, the helical staircase arrangement of subunits can be formed by the ADP-bound helicase alone and does not require a ssDNA substrate. For comparison, we analyzed the helical parameters and pore size of the SVTag•DNA•AMPpNp structure (26). In that complex, the synthetic DNA substrate was present at full occupancy, and the helical z-range was larger (14 Å) with a helical step of 2.9 Å (Table 3). The helical pore size and the average domain rotations were similar to those calculated for fbRep•ADP and pcRep•ADP. The HPV-E1•ADP•DNA complex (25) also shows similar parameters, although for both the SVTag and E1 complexes the domain rotations are more complex (Table 3).

### Cryo-EM structure of a Redondovirus Rep double hexamer

Although hexamers were the most abundant oligomers observed on cryo-EM grids, we also observed larger particles that could be interpreted as two Rep hexamers (Fig. 4A), consistent with the higher mass species observed in SEC-SAXS experiments (Table 1). Because the number of dodecamer particles was smaller than that available for the fbRep hexamer structures, we carried out reconstructions applying C_1_ and C_6_ symmetry to ATPγS-bound fbRep dodecamers (Table 2, Supplemental Figs. 10 & 11). A symmetric reconstruction with 28,346 particles led to a readily interpretable structure with an overall resolution of 2.95 Å. The same reconstruction carried out without symmetry constraints showed an indistinguishable, but lower quality structure at 3.32 Å.

**Figure 4.**
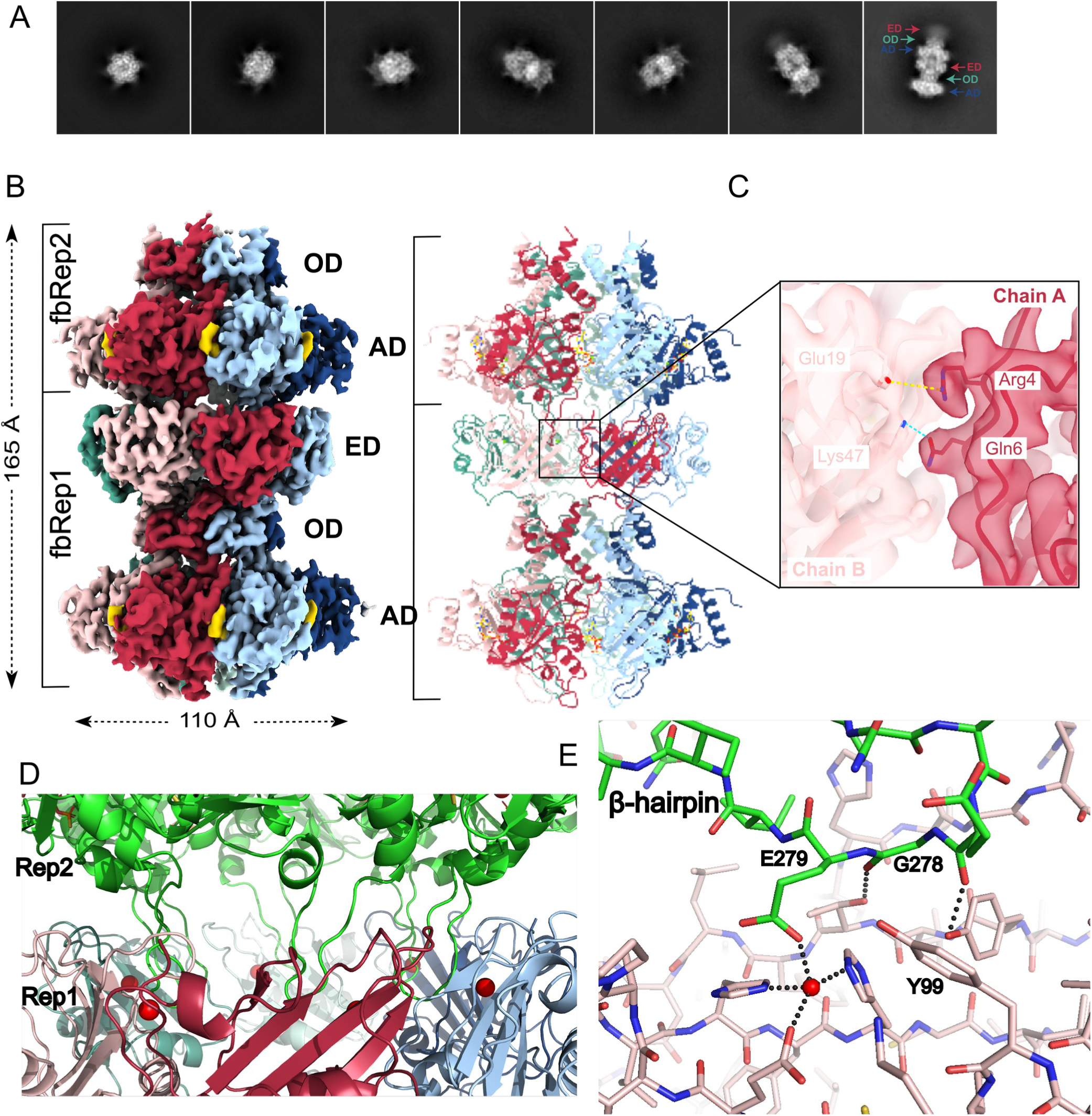
Cryo-EM structure of fbRep•ATPγS double hexamer. A. Representative 2D class averages of fbRep dodecamers. Diffuse density for the endonuclease domains is found only at one end of the particles. B. Reconstruction and cartoon representation of the fbRep dodecamer (C_1_ symmetry). All three domains (ED, OD, AD) are well-defined in fbRep1. The endonuclease domain is poorly ordered in fbRep2, similar to what is seen in the fbRep•ATPγS hexamer. C. Closeup of the interface formed by adjacent endonuclease domains in fbRep1. Few polar or hydrophobic interactions are formed between subunits. D. The distal β-hairpins from fbRep2 (shown in green) interact with the endonuclease domains of fbRep1. E. Closeup of a distal hairpin-ED interaction. Glu279 in the hairpin acts as a phosphate mimic, coordinating magnesium in the endonuclease active site. Gly278 is essential for this interaction because a side chain would be sterically disallowed.

The two hexamers in the fbRep dodecamer associate in a head-to-tail fashion, with a clearly resolved endonuclease domain (ED) ring of one Rep (fbRep1) interacting with the helicase domains of a second Rep (fbRep2, Fig. 4B). The EDs of fbRep2 have diffuse density similar to that observed for isolated fbRep hexamers. Since the ED density for fbRep1 was easily interpretable, we built the domains manually, including well-defined active site magnesium ions. To provide an independent experimental model of the fbRep endonuclease domain, we crystallized it, determined its structure using X-ray diffraction, and refined it to 1.80 Å resolution (Supplemental Table 1, Supplemental Fig. 12). The fbRep1 EDs superposed onto the high resolution ED with r.m.s.d. values of 1.13-1.34 Å across residues 2-109. The ED loops connecting strands β3 and β4 and the loops connecting α3 and β7 adopt different conformations in the fbRep hexamer compared to the isolated domain, explaining the higher than expected r.m.s.d. values.

The interfaces between EDs in the fbRep1 ring lack the extensive contacts normally associated with an oligomeric protein. A salt bridge between Arg4 and Glu19 and a hydrogen bond between Gln6 and Lys47 provide the only polar interactions, and the remainder is comprised of van der Waals contacts with little hydrophobic surface buried (Fig. 4C). The lack of a substantial interface between subunits may explain why the EDs in fbRep hexamers do not form well-defined oligomeric rings, despite being pre-organized to do so by the hexameric helicase domains. The ED hexamer in fbRep1 is instead stabilized through interactions with the distal β-hairpins of the ATPase subdomains from fbRep2 (Fig. 4D). Each β-hairpin binds to the ED β-sheet, with the nuclease active site at the center of the interface (Fig 4E). Two β-hairpin residues appear to play key roles. Glu279 functions as a phosphate mimic, coordinating the active site magnesium ion with its carboxylate side chain, and Gly278 allows a close approach of the β-hairpin to the ED β-sheet. Direct interactions with the peptide backbone of the β-hairpin, together with hydrophobic and polar interactions that could be performed by alternative side chains provide additional, but less specific contacts that flank the active site.

Gly278 is invariant among the redondoviruses and Glu279 is strongly conserved, but replaced by Gln or Asn in some viruses, either of which could coordinate a carbonyl oxygen to the active site magnesium ion in the endonuclease domain interaction. The conservation of Gly278 is particularly compelling because its backbone dihedral angles are within the allowed region for β-strands, implying that it is the lack of a side chain that is most important in this position and that the head-to-tail rep dodecamer is therefore likely to be functionally relevant. Not all CRESS-DNA Rep proteins are expected to have this interaction; the circoviruses and geminiviruses, for example, lack the distal β-hairpin motif. Other viral Reps could, however, form alternative interactions to form head-to-tail hexamer interactions.

We considered two approaches to relate the full-length fbRep structure represented by fbRep1 (Fig. 5A) to isolated Rep hexamers, where the EDs adopt an ensemble of configurations. The first was to calculate predicted scattering profiles for fbRep1 and compare to the hexameric component from SEC-SAXS analysis of the fbRep•ATP complex. However, fbRep•ATP exists primarily in the dodecamer form at the concentrations required for SAXS analysis and the hexameric scattering component is too weak to reliably model. We therefore attempted to model the SAXS profiles from fbRep•ADP and apo-fbRep hexamers (Supplemental Fig. 13). The fit to fbRep•ADP was reasonable (χ^2^=2.1) but the fit to apo-fbRep was poor (χ^2^=4.8), as expected based on the conformational changes that occur upon nucleotide binding (Table 1). The second approach was to dock fbRep1 into the density of the fbRep•ATPγS cryo-EM structure contoured at σ=1. As shown in Fig. 5B, the ED ring occupies the diffuse toroidal density observed in the isolated hexamer. Thus, the arrangement of EDs observed in fbRep1 is likely to be present in the isolated hexamer, but as an ensemble of closely related configurations due to the weak intersubunit contacts that are formed.

**Figure 5.**
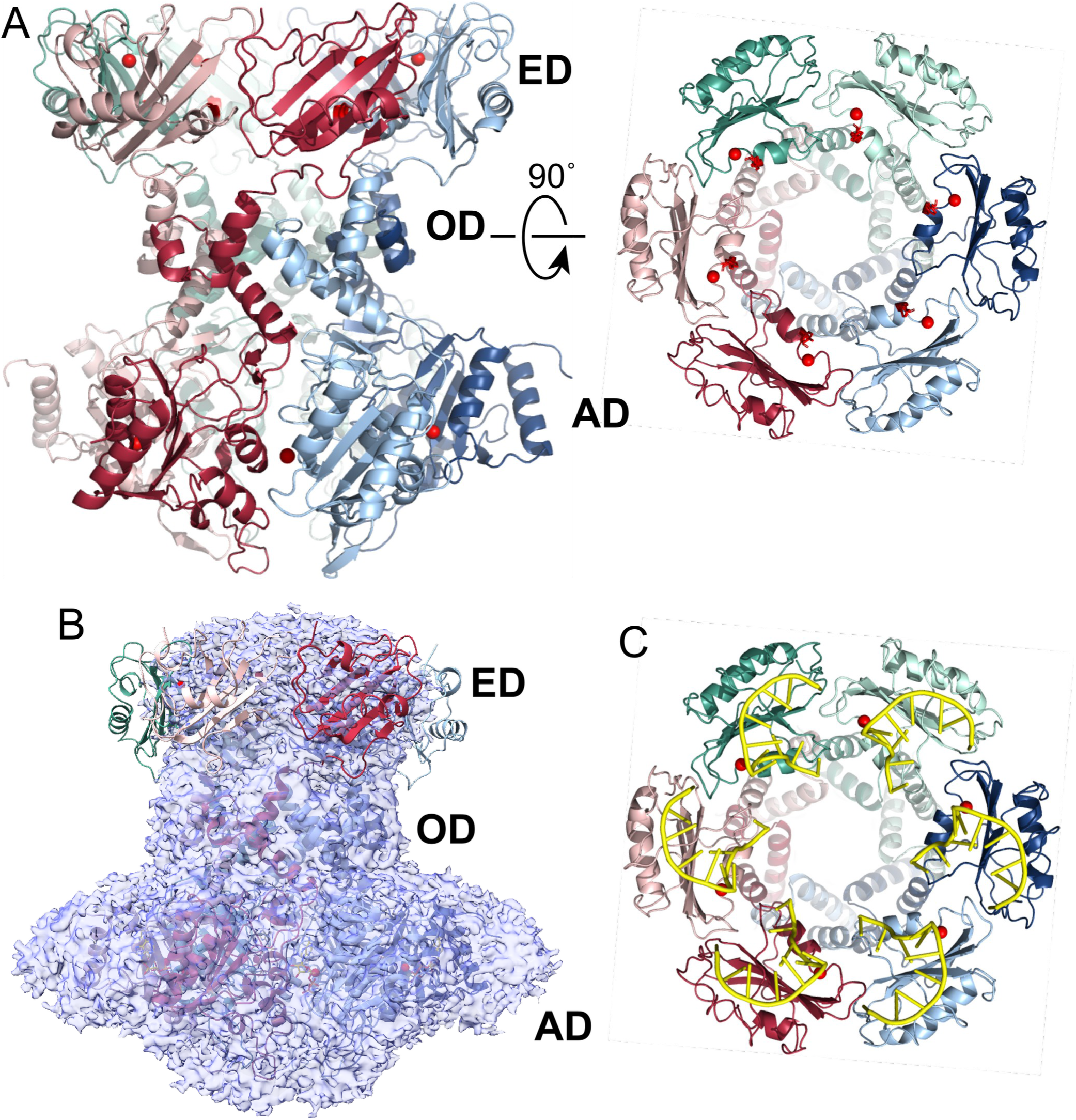
Structure of the Rep hexamer. A. Ribbon representation of Rep as seen in fbRep1 within the fbRep•ATPγS dodecamer. The orthogonal view (right) shows the external face of the endonuclease domains, where a large bowl-shaped cavity is formed by the ED ring. The proximal pore formed by the oligomerization domains is visible at the bottom of the cavity. Magnesium ions are drawn as red spheres. B. Superposition of fbRep1 on the 2.36 Å fbRep•ATPγS hexamer map, contoured at σ=1 to emphasize the diffuse endonuclease domain density. C. Model of the fbRep hexamer with origin ssDNA hairpin bound. The model is based on the PCV2 endonuclease domain-ssDNA complex structure (20).

The cavity formed by the hexameric ED ring is shaped like a large bowl that could accommodate the DNA cruciform substrate that is recognized and cleaved by Rep (Fig. 5A). To verify this, we generated a model of ssDNA-bound fbRep1 using the ED•ssDNA structure reported for pcRep to place the DNA (20). Although the fbRep and pcRep EDs share minimal sequence identity overall, the residues involved in origin recognition and the origin sequences themselves are similar. As shown in Fig. 5C, multiple DNA hairpins can bind to and be cleaved within the bowl-shaped ED cavity as observed in fbRep1, with no steric hindrance. The inward-facing ED active sites should therefore have access to either an external origin sequence, an origin sequence translocating into the Rep helicase channel, or an origin sequence on the non-translocating strand that is separated by the helicase. Since Rep is expected to translocate ssDNA through the oligomerization and ATPase domain pores in the N-terminal to C-terminal direction (26, 32), separation of dsDNA should occur within the ED bowl where both strands can be accessed by ED active sites.

Although the fbRep dodecamer shown in Fig. 4 might be expected to represent two SF3 helicases that work in tandem to translocate along ssDNA, the structure suggests that this is unlikely to be the case. The distal β-hairpins of fbRep2 lock the helicase domains into a symmetric ATP configuration and the domain rotations associated with helicase function upon ATP hydrolysis appear unlikely. Similarly, the endonuclease domains of fbRep1 are inactivated by binding of the distal β-hairpins from fbRep2. Thus, the dodecamer may represent a form of Rep with one functional endonuclease hexamer (on fbRep2) and one functional helicase hexamer (on fbRep1). Assuming that ssDNA can freely translocate through the wider (ATP state) proximal pore of fbRep2, the dodecamer could function as a highly processive form of the Rep enzyme.

## Discussion

CRESS-DNA viruses are ubiquitous on Earth, but the function of their encoded replication initiation proteins —Reps—has been elusive. Here we take advantage of the Redondovirus system to develop multiple views of Rep function, including the first experimental structure of a full-length Rep hexamer. Both biophysical and cryo-EM studies support a hexameric assembly as a major Rep form, but Rep can readily interconvert between pentamers and hexamers, as well as form dodecameric assemblies. Each of these forms may have functional roles.

The cryo-EM structures of ATPγS and ADP-bound fbRep allowed us to compare these states and relate the results to current models for SF3 helicase function. We find that fbRep•ATPγS adopts a symmetric configuration without a helical staircase arrangement of DNA-binding loops. This preference can be explained by optimization of quaternary interactions involving arginine finger motifs interacting with the γ-phosphate of ATPγS. A similar structure was reported for SVTag bound to ATP (27). In the ADP-bound state, fbRep adopts a conformation where the ATPase domains rotate by ∼4° successively around the hexamer, resulting in a helical arrangement of DNA-binding loops and an opening of the helicase distal pore. Similar structures have been observed in other SF3 helicases, but those have been with DNA bound (23, 25, 26). The fbRep•ADP structure indicates that ssDNA interactions are not required for the staircase structure to be formed and that this configuration may be preferred in the ADP-bound state. The SVTag•ADP hexamer crystallized in a nearly symmetric configuration, arguing that both forms are energetically accessible to the helicases.

Current models of SF3 helicase function do not involve all-ATP or all-ADP bound states (25, 26). Reaction intermediates of active SVTag•DNA complexes trapped by freezing on cryo-EM grids indicate that the AB and BC subunit interfaces are ATP-bound (least rotated) and the remaining interfaces have ADP or no nucleotide bound (more rotated) (26). ATP hydrolysis and phosphate release at AB and BC generate ADP sites and ATP binding to DE and FA generate ATP sites, with the associated intersubunit remodeling and domain rotations to generate cyclic permutations of the staircase configuration. These cyclic motions are coupled to unidirectional ssDNA diffusion along the helical ratchet formed by the DNA-binding loops. Since it can be difficult to compare the structural results from different studies, we used simple geometric parameters to describe the helical nature and pore dimension of fbRep and compared the results to other SF3 helicases (Table 3). This quantitative description should be generally useful to other hexameric AAA+ ATPases, provided a reference symmetry axis can be suitably defined. Thus, the fbRep complexes described here provide support for emerging models of SF3 helicase function and establish Redondovirus Rep as a useful model system for studying these enzymes.

The structure of the head-to-tail dodecameric fbRep complex provides new insights into Rep structure and function. Because the endonuclease domains are sequestered into a well-defined hexameric ring, the fbRep dodecamer provides a model for the full-length viral Rep hexamer. Although it was assumed that the EDs would occupy the diffuse density cloud evident in the fbRep hexamer structures described here and in the pcRep•ADP hexamer previously reported (23), the orientations of the domains, the location of the active sites with respect to the helicase pore, and the nature of interactions between subunits were all unknown. Since the fbRep1 hexamer observed in the dodecamer fits well within the isolated fbRep•ATPγS hexamer density, we suggest that fbRep1 represents a functional member of an ensemble of nearly identical structures present in isolated hexamers.

The inward facing active sites of the endonuclease domains may be functionally important in the initiation of replication and in formation of ssDNA genome circles. When Rep initially cleaves the nonanucleotide sequence at the stem-loop viral replication origin, a 5’-phosphotyrosine linkage is formed to the DNA. Once a new copy of the origin is available following polymerase extension of the 3’-end, a second ED can cleave this sequence in the same manner, releasing a 3’-end within the ED ring cavity. The 3’- and 5’- ends of the ssDNA genome would therefore be in close proximity and an intact DNA circle can be formed and released by attack of the original 5’-linkage by the new 3’-end. The endonuclease arrangement we observe could also explain how stem-loop destabilization might occur as described for the ‘RCR melting pot’ model of CRESS-DNA viral replication (33). In the melting pot model, both stem-loops of the cruciform structure formed in the supercoiled genome origin are engaged in a destabilized initiation complex where template switching can occur once Rep nicks the DNA. The six DNA-binding surfaces lining the bowl-shaped ED cavity in fbRep1 might be well suited for this mechanism.

The primary interactions we observe between Rep hexamers in the dodecamer involve solvent-exposed β-hairpins emerging from the distal pore of the helicase and the active sites of the endonuclease domains. Conservation of the key residues involved suggests that dodecamers may play a functional role in redondovirus biology and for many other CRESS-DNA viruses. It is not yet clear what that role may be, but one possibility is that a head-to-tail dodecamer could simply be a more processive version of hexameric Rep. This could be important since we know that Rep subunits readily dissociate in solution. The pentamer-hexamer transition could be used for loading ssDNA into the helicase pore, and the same transition could result in premature dissociation. Formation of the presumably more stable dodecamer on a ssDNA substrate could inhibit dissociation and result in higher processivity. There is precedent for two SF3 hexamers working together on a single DNA substrate. SVTag and HPV-E1 both form head-to-head dimers of hexamers at the replication origin and melt the DNA to initiate bidirectional DNA replication in their eukaryotic hosts (34, 35).

A primary goal of this work has been to use Rep structural models to gain insight into viral replication and DNA packaging. Published models of viral Rep function are mostly schematic in nature and are generalized versions of rolling circle replication from plasmid and related viral systems. The lack of mechanistic models has been partly due to the lack of structural data for CRESS-DNA Reps. Although a number of endonuclease structures have been reported, the only structural data available for hexameric CRESS-DNA Rep helicases has been the recent cryo-EM structure of pcRep bound to ADP. The more distantly related AAV2 Rep has been well-studied, with structural models available, but the oligomeric state, replication mechanism of the virus and nature of the replication origin are all different than that of Redondovirus Rep. The same limitations exist at the biochemical level. Origin recognition and cleavage by Rep have been demonstrated for several CRESS-DNA systems, including fbRep (7). However, demonstration of helicase activity has been more difficult (36) and how that helicase activity is used in replication is unclear.

A simple but important mechanistic question is which DNA strand is translocated by Rep (Fig. 6). If Rep engages the (+) strand that it has just cleaved, a dead-end complex is formed on the 3’-side of the nick, since Rep would be bound at the attached 5’-end, with nothing to translocate (Fig. 6A, top). If Rep engages the (+) strand on the 5’-side of the break, translocation could occur in the opposite direction from replication, and it is not clear how this would be productive (Fig. 6A, bottom). A more plausible mechanism (Fig. 6B) is that Rep engages the (−) strand, perhaps the destabilized (−) strand stem-loop formed during origin recognition (33). Rep could then move ahead of the replication fork and strand separation would occur within the endonuclease cavity. When a newly synthesized (+) strand origin becomes available, it could be recognized and cleaved by one of the five untethered EDs and a new genome circle can be formed by trans-esterification at the original 5’-linkage. Experiments to test this mechanism and alternative mechanisms can be more readily considered, designed and modeled with the structures reported here.

**Figure 6.**
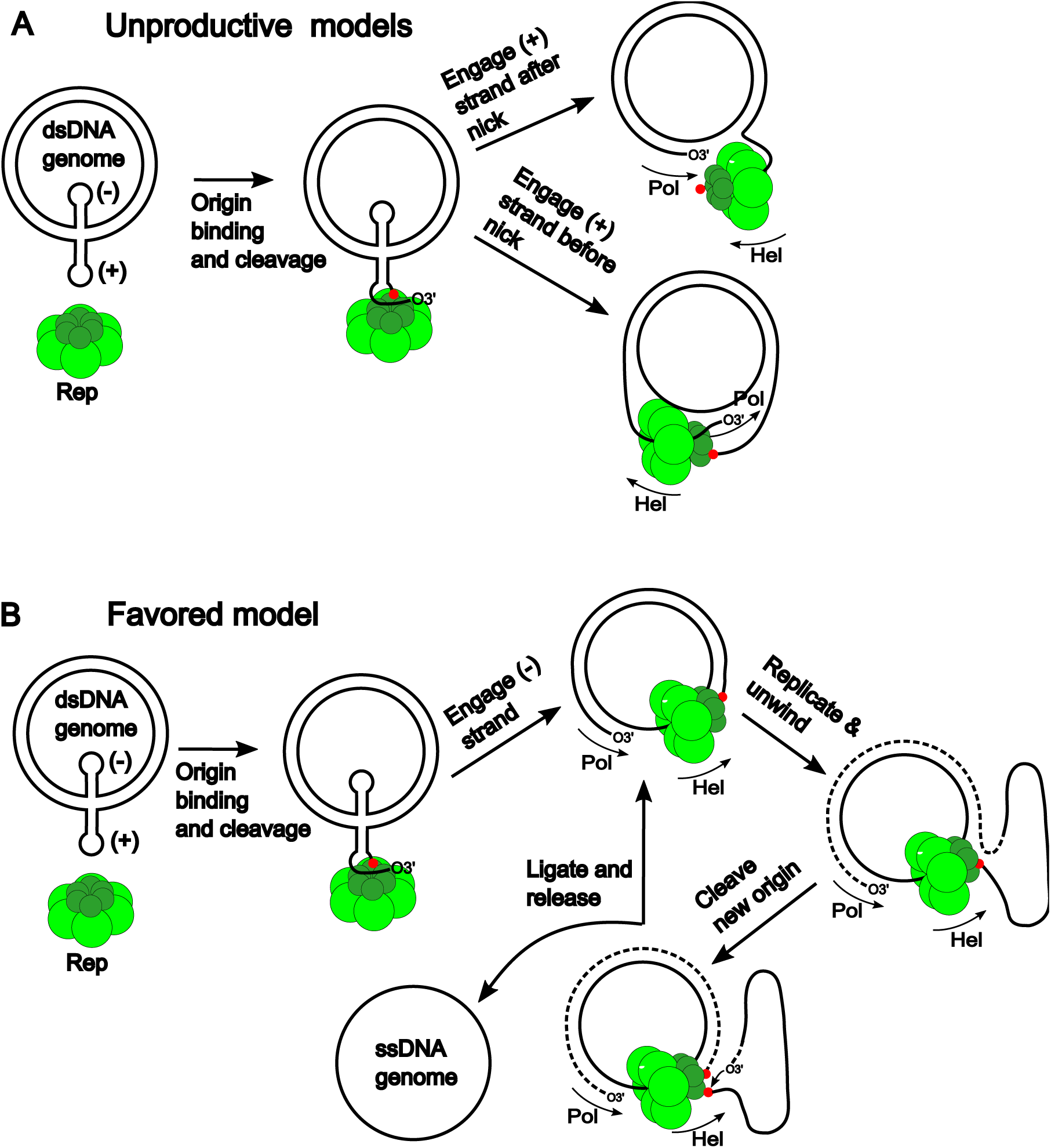
Possible mechanisms of Rep-assisted viral replication. (A) Rep recognizes the (+)-strand origin in the dsDNA viral genome that has been generated and supercoiled by the host. The (−)-strand may also be engaged by the Rep hexamer as suggested (33) but is not shown here. Rep cleaves the ssDNA to form a covalent 5-phosphotyrosine linkage and release a free 3’-end that is extended by host polymerase. Rep could load on the (+)-strand after the nick (top view), resulting in a helicase dead-end. Loading onto the (+)-strand before the nick results in opposing helicase and replication directions. (B) If Rep loads onto the (−)-strand, the DNA can be unwound to provide (+)-strand ssDNA. When a new origin sequence is replicated, Rep can cleave it and catalyze ligation of the extruded DNA to produce ssDNA viral genomes. Rep is represented by a hexamer here for simplicity, but the functional form could be a dodecamer. 5’-covalent linkages to Rep are indicated by red spheres. The direction of polymerase extension and Rep translocation are indicated by arrows.

There are several limitations of the work described here. It would be valuable to complement the structural studies with further functional studies and efforts are underway on several fronts. It has not yet been possible to grow Redondoviruses experimentally in their natural amoeba hosts, but in the future, it would be valuable to generate predictions from in vitro data and confirm them with in vivo tests. Developing a robust in vitro helicase assay, perhaps coupled to origin cleavage and replication, would also be important to test mechanistic models. We have so far been unable to demonstrate helicase activity with Redondovirus Rep protein using standard flap and fork substrates. Since helicase loading may be coupled to origin recognition and cleavage, more specific substrates may be required that have the sequences and properties of native viral DNAs.

In summary, we present the first high resolution structures for Redondovirus Rep proteins, which provide a tractable model for studies of Rep proteins broadly. We report that conformational changes associated with ATP hydrolysis in fbRep strongly support models of SF3 helicase function and the structure of full-length fbRep and dodecameric fbRep provide key insights into Rep function. Going forward, it will be valuable to generate structures for additional functional forms, such as Rep covalently bound to DNA, and develop diverse in vitro assays of function.

## Methods

### Expression and purification of Rep

The DNA sequence of Vientovirus FB Rep (fbRep) (GenBank MK059763.1) was codon-optimized for bacterial expression and cloned into a pCDFDuet expression vector (Novagen) containing an N-terminal His_7_-Flag-SUMO tag. The fbRep endonuclease domain 2-114 (fbRep-ED) was amplified by PCR and cloned into the same vector. Full-length fbRep and fbRep-ED were expressed in HI-Control BL21(DE3) (Lucigen) using 2xYT medium supplemented with 2 mM MgSO_4_ and 50 μg/mL streptomycin at 37°C until an optical density at 600 nm of 1.8-2.2 was reached. The temperature was reduced to 20°C and expression was induced with 0.1 mM isopropyl-β-d-1-thiogalactopyranoside (IPTG) for 5 hours. Cells were harvested by centrifugation and frozen at −80°C.

Cell pellets of fbRep were resuspended in lysis buffer (20 mM NaKPO4 pH 7, 500 mM NaCl, 10 mM imidazole, 10mM β-mercaptoethanol (β-ME), 1X Protease Inhibitor) and lysed mechanically using an Emulsiflex-C3 high pressure homogenizer (Avestin, Inc). The clarified supernatant was loaded onto 5 mL of Ni-NTA Superflow Resin (Qiagen) equilibrated with 10 column volumes of lysis buffer. Ni-bound Rep was washed with 10 column volumes of 20 mM NaKPO4 pH 7, 500 mM NaCl, 20 mM imidazole, 10 mM β-ME and then eluted in 30 mL of 20 mM NaKPO4 pH 7, 500 mM NaCl, 250 mM imidazole, 10 mM β-ME. The His_7_-Flag-Sumo affinity tag was cleaved by ULP1 protease (Life Sensors) while dialyzing against 20 mM Tris-HCL pH 7.4, 500 mM NaCl, 10 mM β-ME. The cleaved affinity tag was removed from solution by a second Ni-NTA column. Rep was further purified by anion exchange using a Source™ 15Q column (GE Healthcare) with a sodium chloride gradient and size exclusion chromatography using Superdex 200 (GE Healthcare) and Superose 6 Increase 10/300 GL (GE Healthcare) columns at room temperature in 20 mM HEPES–NaOH pH 7.5, 300 mM NaCl, 0.1mM TCEP.

fbRep-ED was purified using Ni-NTA chromatography and Ulp1 cleavage as described, except that chromatography buffers contained 300 mM NaCl, and the dialysis buffer contained 100 mM NaCl. fbRep-ED was further purified by cation exchange chromatography using heparin Sepharose (Cytiva) with a NaCl gradient at pH 7.4 and a Ni-NTA flow-through step to remove uncleaved fusion protein. fbRep-ED was further purified by size-exclusion chromatography using a Superose 6 Increase 10/300 GL column (GE Healthcare) at room temperature in 20 mM HEPES pH 7.5, 150 mM NaCl, 0.1mM TCEP.

Proteins were concentrated using Millipore concentrators, concentrations determined using UV absorption at 280 nm, and flash-frozen in liquid nitrogen after addition of glycerol to 10%. Aliquots were stored at - 80°C.

### Size-exclusion Chromatography in line with Multi-angle Light Scattering (SEC-MALS)

SEC-MALS experiments were performed on 3-6 mg/mL full-length Rep protein with a Superose 6 10/300 GL column (GE Healthcare) at 0.5 ml/min at room temperature in buffer containing 20 mm HEPES-NaOH pH 7.5, 300 mm NaCl or 1 M NaCl, and 0.1mM TCEP. Absolute molar mass of fbRep was determined in-line using multi-angle light scattering from the column eluent at 18 angles using a DAWN-HELEOS II MALS detector (Waters/Wyatt Technology Corp., Santa Barbara, CA, USA) operating at 658 nm. Protein concentration of the eluent was determined using an in-line Optilab T-rEX Interferometric Refractometer (Waters/Wyatt Technology Corp.). The weight-averaged molar masses of species within defined chromatographic peaks were calculated using the ASTRA software version 8.2.1 (Waters/Wyatt Technology Corp.), by construction of Debye plots (KC/Rθ versus sin^2^[θ/2]) at one second data intervals. The weight-averaged molar mass was then calculated at each point of the chromatographic trace from the Debye plot intercept, and an overall average molar mass was calculated by weighted averaging across the peak.

### Sedimentation Velocity Analytical Ultracentrifugation (SV-AUC)

SV-AUC experiments were performed at 20°C with an Optima AUC analytical ultracentrifuge (Beckman Coulter) and a TiAn50 rotor with two-channel epon charcoal-filled centrepieces and sapphire windows. In SV-AUC analysis of the full-length fbRep in different nucleotide states, 400 μL of 8.7 μM fbRep protein samples (A_280_: 0.6, 0.35 mg/ml) with and without 5 mM MgCl_2_ and 5 mM ATP-γ-S (or ATP, or ADP) were prepared in a buffer containing 20 mM HEPES-NaOH pH 7.5, 300 mM NaCl and 0.1 mM TCEP. SV-AUC analyses of fbRep at different salt concentration were performed in a buffer containing 20 mM HEPES-NaOH pH 7.5, 1 mM DTT and salt concentration varying from 250 mM to 1 M. Sedimentation velocity profiles were collected every 30 s for 200 boundaries at 40,000 rpm. Data were fit using the Lamm equation model, as implemented in the program SEDFIT (37). After optimizing meniscus position and fitting limits, the sedimentation coefficients (*s*) and best-fit frictional ratio (*f/f_0_*) were determined by iterative least squares analysis. Sedimentation coefficients were corrected to S_20,w_ based on the calculated solvent density (ρ) and viscosity (η) derived from chemical composition by the program SEDNTERP (38). Figures were prepared using the program GUSSI (39)

### Size-exclusion chromatography in line with synchrotron small-angle X-ray scattering (SEC-SAXS)

SEC*-*SAXS data were collected at beamline 16-ID (LiX) of the National Synchrotron Light Source (NSLS) II at Brookhaven National Laboratory (Upton, NY) (40) and the BioSAXS beamline ID7A1 at Cornell MacCHESS (41) using a Superdex 200 5/100 column (GE Healthcare). At both beamlines, the BioXTAS RAW software (Version 2.1.1) (42) was used for data processing and preliminary analysis. All measurements were performed in 20 mM HEPES-NaOH pH 7.5, 300 mM NaCl, 0.1 mM TCEP with or without 1 mM MgCl_2_ and 1mM nucleotide (or ATP or ADP). Samples were injected at 3-5 mg/mL of fbRep protein in 100 µL volumes at room temperature. Radiation damage was not observed in the samples. A summary of the SAXS parameters is given in Table 1.

At beamline 16-ID, data were collected at a wavelength of 1.0 Å in a three-camera conformation, yielding accessible scattering angle with 0.005 < q < 3.0 Å^−1^, where q is the momentum transfer, defined as q = 4π sin(θ)/ λ, where λ is the X-ray wavelength and 2θ is the scattering angle. The data to q<0.5 Å^−1^ were used in subsequent analyses. Samples were loaded into a 1-mm capillary for ten 1-s X-ray exposures. At ID7A1, data were collected at a wavelength of 1.1 Å in with an Eiger 4.0 detector, yielding accessible scattering angle with 0.008 < q < 0.67 Å^−1^. A series of 707 images for fbRep only, 500 images each for ADP-bound and ATP-bound fbRep were collected, with 2-s exposures per image.

### SAXS Analysis

SVD-EFA analyses of the SEC-SAXS datasets were performed as implemented in the RAW program (31). Buffer subtracted profiles were analysed by singular value decomposition (SVD) and the ranges of overlapping peak data were determined using evolving factor analysis (EFA). The determined peak windows were used to identify the basis vectors for each component and the corresponding SAXS profiles were calculated. When fitting manually, the maximum diameter of the particle (D_max_) was incrementally adjusted in GNOM (43) to maximize the goodness-of-fit parameter, to minimize the discrepancy between the fit and the experimental data, and to optimize the visual qualities of the distribution profile. DENSS (44) was used to calculate the *ab initio* electron density directly from GNOM outputs, as implemented in the program RAW. For each nucleotide state, twenty reconstructions of electron density were performed in the slow mode with default parameters and subsequently averaged and refined with or without C6 symmetry restraints. Reconstructions were visualized using either UCSF ChimeraX version 1.9 (45) or PyMOL version 3.1 (Schrodinger, LLC, New York, NY, USA).

Hybrid bead-atomistic modeling of hexameric fbRep was performed using the program CORAL (46), where the known structure was fixed in composition and inventory not resolved by X-ray crystallography was modeled as coarse-grain beads. Ten independent calculations for each protein were performed and yielded comparable results. The final models were assessed using the program FoxS (47). The models were rendered using the program PYMOL.

### Mass Photometry

Mass photometry experiments were conducted using a Refeyn TwoMP instrument (Refeyn Ltd., Littlemore, Oxford, UK). Prior to the experiment, a standard curve relating particle contrasts to molecular weight was constructed with molecular weight standards made with 3 nM thyroglobulin and 10 nM β-amylase. Samples at 20 μL volumes were evaluated in reaction chambers formed by placing silicone Sample Well Cassettes with 6-3.0 mm diameter, 1.0 mm depth, and 11mm × 15 mm OD onto MassGlass UC coverslips (Refeyn Ltd.). Mass photometry signals were recorded for 60 seconds at room temperature. Data acquisition and analysis were performed using AcquireMP and DiscoverMP (Refeyn Ltd., version 2022 R1). Experiments using 100 nM purified fbRep were conducted in a buffer consisting of 20 mM HEPES-NaOH pH 7.5, 300 mM NaCl and 0.1 mM TCEP. Additionally, experiments were performed in the presence or absence of 5 mM MgCl_2_ and 5 mM ATPγS or ADP.

### Cryo-EM sample preparation and data collection

ATPγS-bound and ADP-bound fbRep samples were prepared by incubating 5 μM fbRep protein with 5 mM MgCl_2_ and 5 mM ATP-γ-S (or ADP) in 20 mM HEPES-NaOH pH 7.5, 300mM NaCl, 0.1 mM TCEP and 2.5% glycerol for 30 minutes on ice. Quantifoil^®^ 1.2/1.3 300 mesh Cu (Electron Microscopy Sciences) grids were glow-discharged for 30 seconds at 10 mA using a PELCO easiGlow™ glow-discharge system. For grid samples preparation, a 3 μl protein sample was spotted, blotted for 6s at 4°C with 0 blot force in 100% humidity and plunged frozen into liquid ethane using the Vitrobot Mark IV (Thermo Fisher Scientific). Grids were then stored in liquid nitrogen until screening and data collection. Initial screening and data collection were conducted at the Beckman Center for Cryo-Electron Microscopy at the University of Pennsylvania Perelman School of Medicine (RRID: SCR_022375) on Titan Krios G3i 300 kV electron microscope equipped with a Gatan K3 direct electron detector. For the ATPγS-bound fbRep, 5605 movies were collected with EPU (Thermo Fisher Scientific) using aberration-free image shift (AFIS) strategy at 105,000X magnification with a pixel size of 0.42 Å/pixel. For the ADP-bound fbRep, 3790 movies were collected with EPU using AFIS strategy at 105,000X magnification with a pixel size of 0.43 Å/pixel. Movies were collected as 35 frames in super resolution mode with a defocus range from −1.0 to −3.0 μm and a total exposure of 41.3 e^−^/Å^2^ for ATP-γ-S-bound fbRep dataset and 42.2 e^−^/Å^2^ for ADP-bound fbRep dataset.

### Cryo-EM image processing

#### ATPγS-bound fbRep

5605 movies were motion corrected with binning by a factor of 2 to 0.84 Å/pix using Relion v5.0-beta (48, 49) implementation of MotionCor2-like algorithm (50) followed by estimation of contrast transfer function (CTF) parameters using CTFFIND-4.1.14 (51). The motion corrected micrographs were then imported into CryoSPARC (v4.5.3) (52) and the CTF parameters were estimated using Patch CTF estimation. Micrographs were manually curated to remove low-quality micrographs, and the remaining 5544 micrographs were denoised using Topaz denoising (53) tool prior to particle picking. Initial sets of particles were blob picked from 200 denoised micrographs to obtain templates for full dataset particle picking and initial models for heterogeneous refinement. Initial 98,258 particles were binned by 4 to a box size of 72 pixels during extraction and subjected to a round of 2D classification. Selected 2D classes were re-extracted with binning by 2 to a box size of 144 pixels and subjected to another round of 2D classification to remove obvious junk particles. The remaining 54,440 particles were used in reconstruction of ab initio models (*N=4*), generating two junk models and an intact hexameric and pentameric models. Four decoy models reconstructed from 90 particles were also obtained from the remaining particle stack using *ab initio* reconstruction. Both hexameric and pentameric models were then aligned to C6 symmetry axis and subjected to non-uniform refinement (54). Well-resolved 2D classes from the hexameric and pentameric particle datasets were then used as templates for full dataset template-based particle picking.

5,286,720 template-based picked particles were extracted with binning by 4 at a box size of 72 pixels. The extracted particles were subjected to a round of heterogeneous refinement using a hexamer, a pentamer, and four decoy models resulting into classification of 1,454,002 and 817,824 particles into the hexamer and pentamer models, respectively. The hexameric and pentameric particle subsets were then separately subjected to three rounds of heterogeneous refinement using the same initial models. In each round of heterogeneous refinement for the hexameric and pentameric particle subsets, the particles that were classified into the hexameric or pentameric model were combined and used as the input particles for the next round of heterogeneous refinement while all particles classified into the decoy models were discarded. After the third round of heterogeneous refinement, the combined 1,446,088 hexameric and pentameric particles were extracted with binning by a factor of 2 at a box size of 144 pixels and subjected to several rounds of heterogeneous refinement using the same strategy as previously done until less than 2% of particles are classified into decoy models. The hexameric and pentameric particles from the last rounds of heterogeneous refinement were further sorted with 2D classifications, keeping only particles from well-resolved 2D classes. The hexameric and the pentameric particles from selected 2D class averages were separately subjected to extra rounds of heterogeneous refinement using hexameric or pentameric models with four decoys to ensure clean particle stacks. The final hexameric and pentameric particles were classified into 2D classes to visually assess particle orientations and remove any remaining particles sorted into poorly resolved classes. The 963,513 hexameric and 202,611 pentameric particle stacks were extracted without binning, refined using non-uniform refinement, and exported to Relion for further analysis.

In Relion, the final particle stacks were reextracted without binning to a box size of 288 pixels and then subjected to 3D auto-refinement using the volume and mask from non-uniform refinements performed in CryoSPARC. After post-processing, particles were subjected to CTF refinements followed by Bayesian polishing. After 3D auto-refinement and post processing, the particles were subjected to another round of CTF refinements and Bayesian polishing. 3D refinement of polished hexameric and pentameric particles generated 2.35 Å and 3.22 Å resolution maps, respectively.

To analyze for structural heterogeneity in the data, the polished hexameric particles were subjected to unmasked 3D classification with exhaustive angular searches at a sampling rate of 7.5 degrees (*T=4, K=6*). 3D classification generated a junk class of 418 particles, a stacked double hexamer of 5,955 particles, a poorly aligned hexamer of 220,094 particles, a well-aligned hexamer with low-resolution features consisting of 75,338 particles, and two well-aligned hexamers with high-resolution features consisting of 314,908 and 346,800 particles. The stacked double hexamer of 5,955 particles was used for further analysis of double hexameric particles in the dataset. 3D refinement of the well-aligned hexamer of 75,338 particles generated a 3.18 Å resolution map. The other two highly similar well-aligned hexamers were combined and subjected to a 3D refinement. Refinement of the combined hexamers with 661,708 particles generated a final map of 2.36 Å resolution. Local resolution of all final reconstructed maps was estimated in CryoSPARC at 0.143 threshold.

#### ATPγS-bound Rep double hexamer

The stacked double hexamer from unmasked 3D classification with exhaustive angular searches refined to a resolution of 3.78 Å with C_1_ symmetry and 3.27 Å with C_6_ symmetry. The dataset used in the reconstruction of the double hexamer map consists only of 5,955 particles. To determine if there are more double hexameric particles present in the micrographs, the double hexameric particles were picked from 5544 micrographs using Topaz picking algorithm in CryoSPARC (55). First, the 5955 particles were exported to CryoSPARC, manually re-picked, extracted with binning by 4 at a box size of 96 pixels, and then used to generate a Topaz picking model. Topaz picking on 5544 micrographs obtained 138,554 particles which were then extracted with binning by 4 a box size of 100 pixels and then subjected to a 2D classification. Well-resolved 2D class averages consisting of 29,474 double hexameric particles were then used to generate an *ab initio* model. The model was aligned to C6 symmetry axis and subjected to non-uniform refinement. A round of heterogeneous refinement using the refined model and three decoy models was done to remove junk particles. The remaining 28,462 particles were extracted without binning to a box size of 440 pixels and were used to generate an ab initio model. The ab initio model was then aligned to C6 symmetry axis and subjected to a non-uniform refinement, generating a 3.71 Å resolution map. To obtain a higher resolution reconstruction, the particles were then exported to Relion for further processing.

In Relion, the double hexameric particles were re-extracted and subjected to 3D auto-refinement using the volume and mask generated from non-uniform refinement in CryoSPARC. After post-processing, the particles were subjected to CTF refinements followed by Bayesian polishing. 3D refinement of the polished particles improved the resolution to 3.52 Å. To identify inconsistent features in the refined particle dataset, the particles were subjected to masked 3D classification without image alignment (*T=4, K=6*), resulting to a removal of 116 particles. Refinement of the remaining 28,346 particles improved the resolution to 3.42 Å. The final particles were then subjected to another round of CTF refinements followed by Bayesian polishing. 3D auto-refinement of the polished particles with C_1_ symmetry generated a final model with an overall resolution of 3.32 Å. Imposing C_6_ symmetry on 3D auto-refinement improved the resolution of the map to 2.95 Å. Local resolution of all final reconstructed maps was estimated in CryoSPARC at 0.143 threshold.

#### ADP-bound Rep

Data analysis for 3790 movies was also done in Relion v5.0-beta and CryoSPARC v4.5.3. Micrographs were motion corrected and CTF estimated in the same way as in ATPγS-bound fbRep dataset. Manual curation of 3790 patch CTF-estimated micrographs eliminated 150 suboptimal micrographs, leaving 3640 micrographs for data analysis. Initial 221,667 blob-picked particles from 500 denoised micrographs were extracted with binning by 4 and subjected to a round of 2D classification. Well-resolved 2D classes consisting of 31,160 particles were re-extracted with binning by 2 and subjected to a round of 2D classification. Well-resolved 2D classes consisting of 26,788 particles were then used as templates in template-based picking of particles from whole dataset. 4,802,310 template-picked particles were extracted with binning by 4 at a box size of 72 pixels and subjected to a round of 2D classification. Selected 2D classes consisting of 341,693 particles were then re-extracted with binning by 2 at a box size of 144 pixels and subjected to another round of 2D classification. Selected side and tilted views were subjected to a round of 2D classification, generating well-resolved side and tilted views of 3,093 particles that were used to train a Topaz picking model. 250,032 Topaz picked particles were extracted by binning by factor of 2 at a box size of 144 pixels and subjected to two rounds of 2D classification. 2D class averages of side and tilted views consisting of 8,144 particles were selected for another round of Topaz training. 195,026 Topaz-picked particles were extracted with binning by 2 at a box size of 144 pixels and subjected to a round of 2D classification. Well-resolved 2D classes of tilted and side views of 101,013 particles were then selected and combined with the 341,693 particles. An initial model consisting of 30,000 particles from the combined particle sets were obtained and refined using nonuniform refinement with symmetry relaxation using maximization method. To remove junk particles, several rounds of hetero refinement were performed using the nonuniform refined model and five decoy models. 417,686 particles from the good class were re-extracted without binning at a box size of 288 pixels and subjected to a round of hetero refinement with decoys. 417,679 particles from the good class were then subjected to a round of 2D classification. Well-resolved 2D classes consisting of 342,764 were then selected and used to obtain an *ab initio* model. Nonuniform refinement using the *ab initio* model aligned to C_6_ symmetry axis generated a 3.59 Å resolution map. To obtain an improved reconstruction, the clean particle stack was exported to Relion for particle polishing and further analysis.

In Relion, the clean particle stack was re-extracted and subjected to a 3D auto-refinement and post processing. After post processing, the particles were then subjected to CTF refinements followed by Bayesian polishing. After 3D auto-refinement, the polished particles were subjected to a masked 3D classification without image alignment (*T=4, K=6*), resulting to one good class consisting of 312,272 particles and two bad classes of 12,665 and 17,827 particles. 3D auto-refinement of the 312,272 particles resulted to a 3.39 Å resolution map which were then subjected to CTF refinements followed by Bayesian polishing. 3D auto-refinement of the polished particles at C_1_ symmetry resulted to a final map with an overall resolution of 3.20 Å. Local resolution of the final reconstructed map was estimated in CryoSPARC at 0.143 threshold.

### Cryo-EM model building and refinement

Prior to model building, the final refined maps were first sharpened with phenix.auto_sharpen (56) implemented in Phenix software package version 1.21-5207-000 (57) using b_iso_to_d_cut sharpening method. Molecular models were built using UCSF ChimeraX version 1.9 and Coot version 0.9.8.96 EL (58) and refined with Phenix version 1.21-5207-000. For modeling of 2.36 Å fbRep•ATPγS hexamer, the hexameric model of fbRep generated using AlphaFold Multimer (59) hosted on COSMIC^2^ cloud platform (60) was used to build the oligomerization and ATPase domains. Initially, the truncated AlphaFold multimer model (120–349) was fitted into the map in ChimeraX. The fitted model was manually adjusted to the map in COOT, truncated to 121-327 amino acids, and then refined with phenix.real_space_refine (61). Iterative model building and refinements were done with ligands and water molecules manually added and refined during the final cycles of refinement. The models for 3.18 Å fbRep•ATPγS hexamer and 3.20 Å fbRep•ADP hexamer maps were built and refined as previously described but using the refined 2.36 Å fbRep•ATPγS hexamer without ligands as the initial model. The stacked double hexamer models were initially built using two refined fbRep•ATPγS hexamer models without ligands into the map using ChimeraX. The endonuclease domains were then built into the model residue by residue in Coot. Several cycles of model building and refinement were performed with ligands added during the final cycles of refinement. All refined models were validated against cryo-EM maps using phenix.validation_cryoem (62) and EMRinger (63) implemented in Phenix. A summary of cryo-EM data collection, refinement, and validation statistics can be found in Table 2.

### X-ray Crystallography

Crystallization trials were performed using the hanging drop method by equilibrating a 1:1 mixture of 200 μM fbRep-ED prepared in 20 mM Tris-HCl pH 7.4, 100mM NaCl, 300 μM MgCl_2_, 0.1 mM TCEP and crystallization buffer against the crystallization buffer containing 100 mM ammonium malonate pH 5.8, 200 mM potassium fluoride, 15 % w/v polyethylene glycol 4000, 15% v/v ethylene glycol. Crystals appeared within a week following incubation at 21°C, and the selected crystals were harvested and stored in liquid nitrogen for data collection.

Diffraction data were collected at 100°K and wavelength of 0.92 Å on beamline 17-ID-1 (AMX) at NSLSII (Brookhaven National Laboratory, Upton, NY). Images were processed using the DIALS software package (Diffraction Integration for Advanced Light Sources) (64). Molecular replacement was carried out using Phaser-MR (65) as implemented in Phenix. The structure was solved using an AlphaFold (66) model of the Rep endonuclease domain 2-114 as a search model. The model was iteratively refined in reciprocal space with Phenix and in real space with COOT. Crystallographic data collection and refinement statistics can be found in Supplemental Table 1.

### Structural analysis

Pore channel profiles of ATPγS-bound and ADP-bound fbRep were generated using MOLEonline (67) with cavity parameters of 45 Å probe radius and 3 Å interior threshold, channel parameters of 10 Å origin radius and 20 Å surface cover radius, 1.2 Å bottleneck radius, 3 Å bottleneck tolerance, 1.0 max tunnel similarity, Voronoi scale weight function, starting point at residue 146, and end point at residue 276. Analysis of hydrogen bonding and hydrophobic contacts were performed in Ligplot+ software v2.2 (68) with default parameters. Magnesium ion contacts were calculated in UCSF ChimeraX with center-center distance parameter value of less than or equal 3.10 Å. Structural alignments were also done in UCSF ChimeraX using the matchmaker tool with default parameters. The helical parameters of fbRep and other SF3 family helicases were calculated using locally written programs.

### Map and model visualization

Maps and models were visualized in UCSF ChimeraX version 1.9. Model illustrations were prepared using UCSF ChimeraX version 1.9 or PyMOL version 3.1.

## Acknowledgements

We are grateful to members of the Moiseenkova-Bell, Gupta, Bushman and Van Duyne laboratories for help and suggestions. We acknowledge the use of instruments at the Electron Microscopy Resource Laboratory (EMRL) and at the Beckman Center for Cryo-Electron Microscopy at the University of Pennsylvania Perelman School of Medicine. We are also grateful to former and current staff members of EMRL and Beckman Center for Cryo-EM, especially to Sudheer Molugu for help and training in negative stain and cryo-EM grid sample freezing and Stefan Steimle for assistance in sample grid preparation and data collection on the Titan Krios. We also wish to extend our gratitude to Ronen Marmorstein laboratory especially to Mary Szurgot and Lauren Gardner for assistance with mass photometry data collection. K.G. acknowledges support from the Johnson Research Foundation and NIH Shared Instrumentation Grant S10-OD018483. The SV-AUC, SEC-MALS, and Mass Photometry experiments were performed at the Johnson Foundation Biophysical and Structural Biology Core Facility (University of Pennsylvania, Philadelphia PA). This work is based in part on research conducted at the Center for High-Energy X-ray Sciences (CHEXS), which is supported by the National Science Foundation (BIO, ENG and MPS Directorates) under award DMR-2342336, and the Macromolecular Diffraction at CHESS (MacCHESS) facility, which is supported by award 1-P30-GM124166 from the National Institute of General Medical Sciences, and the National Institutes of Health. The crystallographic data were obtained at the 17-ID-2 beamline, while additional SEC-SAXS data were obtained at 16-ID (LIX) at the National Synchrotron Light Source II, a U.S. Department of Energy (DOE) Office of Science User Facility operated for the DOE Office of Science by Brookhaven National Laboratory under Contract No. DE-SC0012704. This work was supported in part by P30AI045008 (FB) and GM108751 (GV). S. M. was supported by T32-AI007632.

## Data Availability

All cryo-EM maps presented here have been deposited in the Electron Microscopy Data Bank under the following accession codes: EMD-71784 (fbRep•ATPγS, no ED), EMD-71786 (fbRep•ATPγS, with ED), EMD-71788 (fbRep•ATPγS double hexamer, C_1_), EMD-71789 (fbRep•ATPγS double hexamer, C_6_), and EMD-71790 (fbRep•ADP). Model coordinates built into the maps have been deposited in the Protein Data Bank under the following accession codes: 9PQJ (fbRep•ATPγS, no ED), 9PQM (fbRep•ATPγS, with ED), 9PQQ (fbRep•ATPγS double hexamer, C_1_), 9PQR (fbRep•ATPγS double hexamer, C_6_), and 9PQT (fbRep•ADP). Raw micrographs used for the data analysis of ATPγS-bound fbRep and ADP-bound fbRep have been deposited in the EMPIAR database with accession codes EMPIAR-12903 and EMPIAR-12906, respectively. Coordinates and structure factors for the fbRep endonuclease domain have been deposited with accession code 9PQF. SAXS data are available in SASDB under the following accession codes: SASDXP7 (apo fbRep dodecamer), SASDXQ7 (apo fbRep hexamer), SASDXR7 (ATP-bound fbRep dodecamer), SASDXS7 (ADP-bound fbRep dodecamer), and SASDXT7 (ADP-bound fbRep hexamer).

