## Supplemental Data for "Structures of Nucleotide-Bound Redondovirus Rep Protein Link Conformation and Function"

**Supplemental Table 1. Crystallographic data collection and refinement statistics for fbRep-ED 2-114**

| | |
|---|---|
| PDB code | 9PQF |
| X-ray Source | NSLS-II 17-ID-1 |
| Wavelength (Å) | 0.92 |
| Temperature (K) | 100 |
| Detector | DECTRIS EIGER X 9M |
| Resolution range (Å) | 45.13 - 1.8 (1.86 - 1.8) |
| Space group | I 2 2 2 |
| Unit cell (a, b, c, α, β, γ) | 50.944 62.974 64.705 90 90 90 |
| Unit Cell Volume ($Å^3$) | 207585.079 |
| Total reflections | 74196 (7511) |
| Unique reflections | 9971 (965) |
| Multiplicity | 7.4 (7.8) |
| Completeness (%) | 99.98 (100.00) |
| Mean I/sigma(I) | 4.79 (1.00) |
| Wilson B-factor ($Å^2$) | 23.34 |
| R-merge[b] | 0.1938 (0.424) |
| R-meas[c] | 0.2087 (0.4561) |
| R-pim[d] | 0.07636 (0.166) |
| $CC_{1/2}$ | 0.983 (0.891) |
| CC* | 0.996 (0.971) |
| Reflections used in refinement | 9970 (965) |
| Reflections used for R-free | 982 (107) |
| R-work | 0.1799 (0.2460) |
| R-free | 0.2179 (0.2920) |
| CC (work) | 0.969 (0.876) |
| CC (free) | 0.945 (0.869) |
| Number of non-hydrogen atoms | 961 |
| macromolecules | 904 |
| ligands | 8 |
| solvent | 49 |
| Protein residues | 113 |
| RMS (bonds) (Å) | 0.007 |
| RMS (angles) (°) | 0.90 |
| Ramachandran favored (%) | 96.40 |
| Ramachandran allowed (%) | 3.60 |
| Ramachandran outliers (%) | 0.00 |
| Rotamer outliers (%) | 0.98 |
| Clashscore | 1.09 |
| Average B-factor ($Å^2$) | 27.91 |
| macromolecules | 27.54 |
| ligands | 28.89 |
| solvent | 34.51 |

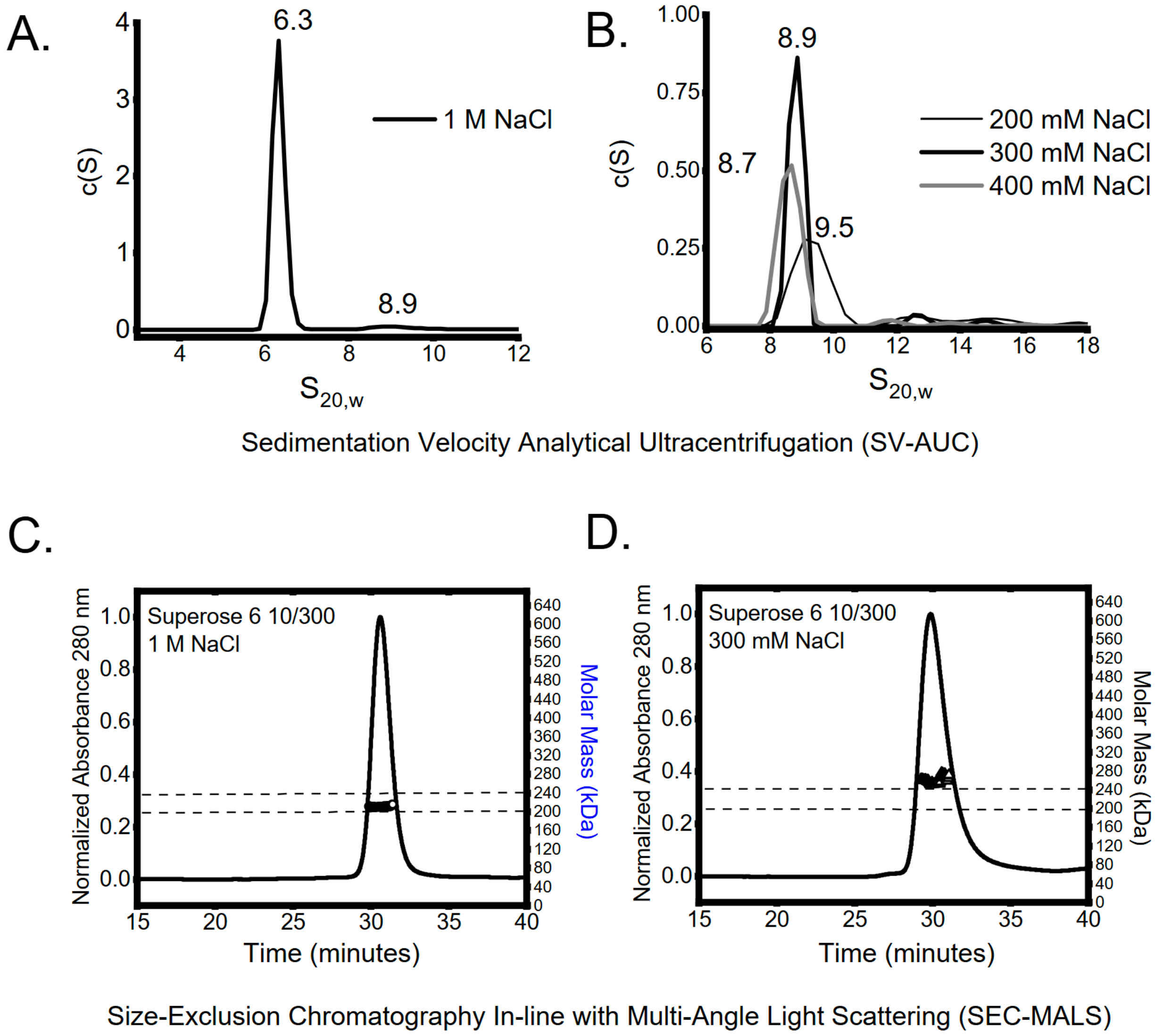


**Supplemental Figure 1. Effects of ionic strength on the oligomeric state of recombinant fbRep. A&B.** Sedimentation Velocity Analytical Ultracentrifugation (SV-AUC) analysis of purified Rep in 1 M NaCl (**A**) or 200-400 mM NaCl (**B**). Shown are c(S) distributions (black lines). In 1 M NaC (**A**), a 6.3S species is most prominent, whereas particles with S values ranging from 8.7-9.5 S are observed, changing in response to increasing ionic strength (**B**). Additional higher order species between 11-16S are also detected in smaller quantities. **C&D.** Size-exclusion chromatography inline with multi-angle light scattering (SEC-MALS) analysis of purified fbRep in 1 M NaCl (**C**) or 300 mM NaCl (**D**). Shown in solid black lines are the SEC elution profiles for fbRep, with absolute weight-averaged molecular mass ($M_w$) (circles) determined by light scattering plotted across the profiles. Dashed horizontal lines denote the masses for pentamers and hexamers, respectively. In 1M NaCl (C), masses most consistent with pentamers are observed (210 kDa ± 1.1), whereas in 300 mM NaCl, masses larger than hexamer are observed (260 kDa ± 6.7).

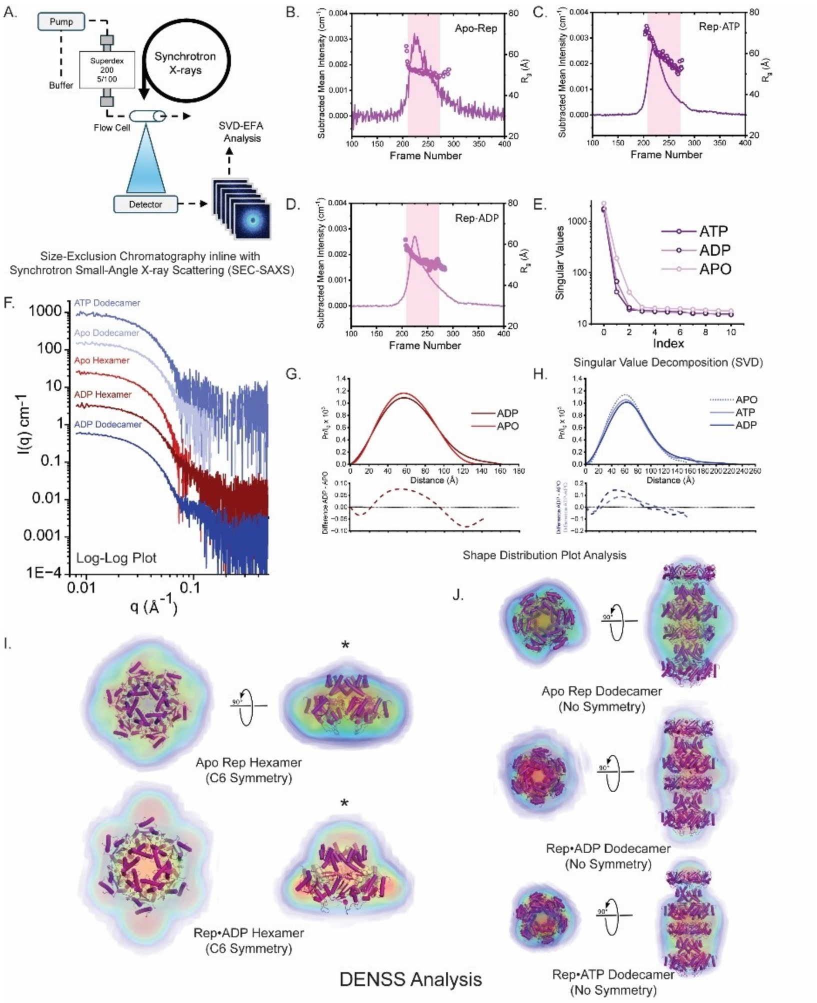


**Supplemental Figure 2. Size-Exclusion Chromatography inline with Synchrotron Small-Angle X-ray Scattering (SEC-SAXS). A.** Experimental Scheme for synchrotron SEC-SAXS analysis is shown. See **Methods** for more details. **B-D.** Lines show integrated mean intensities of X-ray small-angle scattering vs frames recorded during elution of Rep from SEC in its apo form (**B**), and with 1 mM ATP (**C**), or with 1 mM ADP (**D**) present in the running buffer. Circles show the derived radii of gyration ($R_g$) from the background subtracted X-ray scattered profiles. **E.** Singular value decomposition (SVD) analysis of the exposures denoted by pink shading in panels B-D. In each of the three samples tested, the peak region data was best described by three significant species, comprised of dodecamers, hexamers, and aggregates or buffer artifacts. **F.** SAXS profiles for hexamers and dodecamers derived from SVD-EFA analysis (1, 2) of SEC-SAXS data for Rep protein, scaled arbitrarily along the y-axis and shown as a log-log plot. **G&H.** Shape distribution (P(r)) function analysis for hexamers (G) and dodecamers (H), performed using the program GNOM. Parameters derived from this analysis are provided in **Table 1**. In the respective lower panels differences in Pr are shown, illustrating the reorganization of interatomic vectors in response to nucleotide binding of the respective oligomer. **I&J.** Shown is DENSS (3) analysis of the synchrotron SAXS data**.** In orthogonal views for each of the hexamers and dodecamers isolated by SVD-EFA SAXS are *ab initio* electron density reconstructions, docked with the corresponding experimental hexamer models determined (**I**) or with head-to-tail dodecamer models (**J**). Asterisks in Panel I denote spatial discrepancies between the experimental SAXS volumes and where N-terminal endonuclease domains would be expected to reside. Electron density is colored with five contour levels of density rendered with these respective colors: 15σ (red), 10σ (green), 5σ (cyan), 2.5σ (blue), and -0.7σ (blue). The sigma (σ) level denotes the standard deviation above the average electron density value of the generated volume. DENSS figures were generated using the program PyMOL.

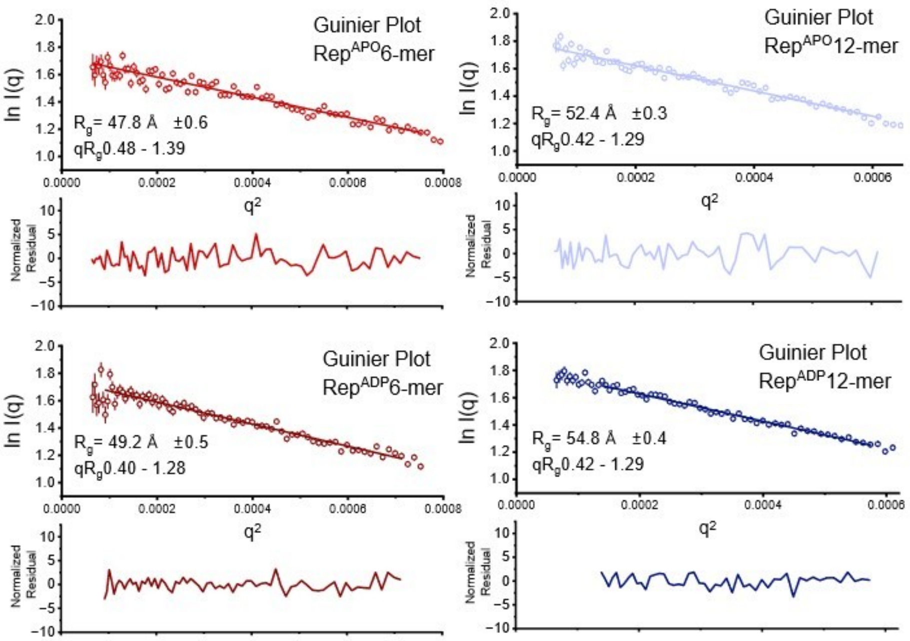


**Supplemental Figure 3. Guinier plot analyses** (lnI($q$) vs. $q^2$) of SAXS data (dots) for Rep oligomers, with residuals from the fitted lines shown below. Monodispersity is evidenced by linearity in the Guinier region of the scattering data and agreement of the $I_0$ and $R_g$ values determined with inverse Fourier transform analysis by the programs GNOM (4) (**Table 1**). Guinier analyses were performed where $qR_g \leq 1.41$.

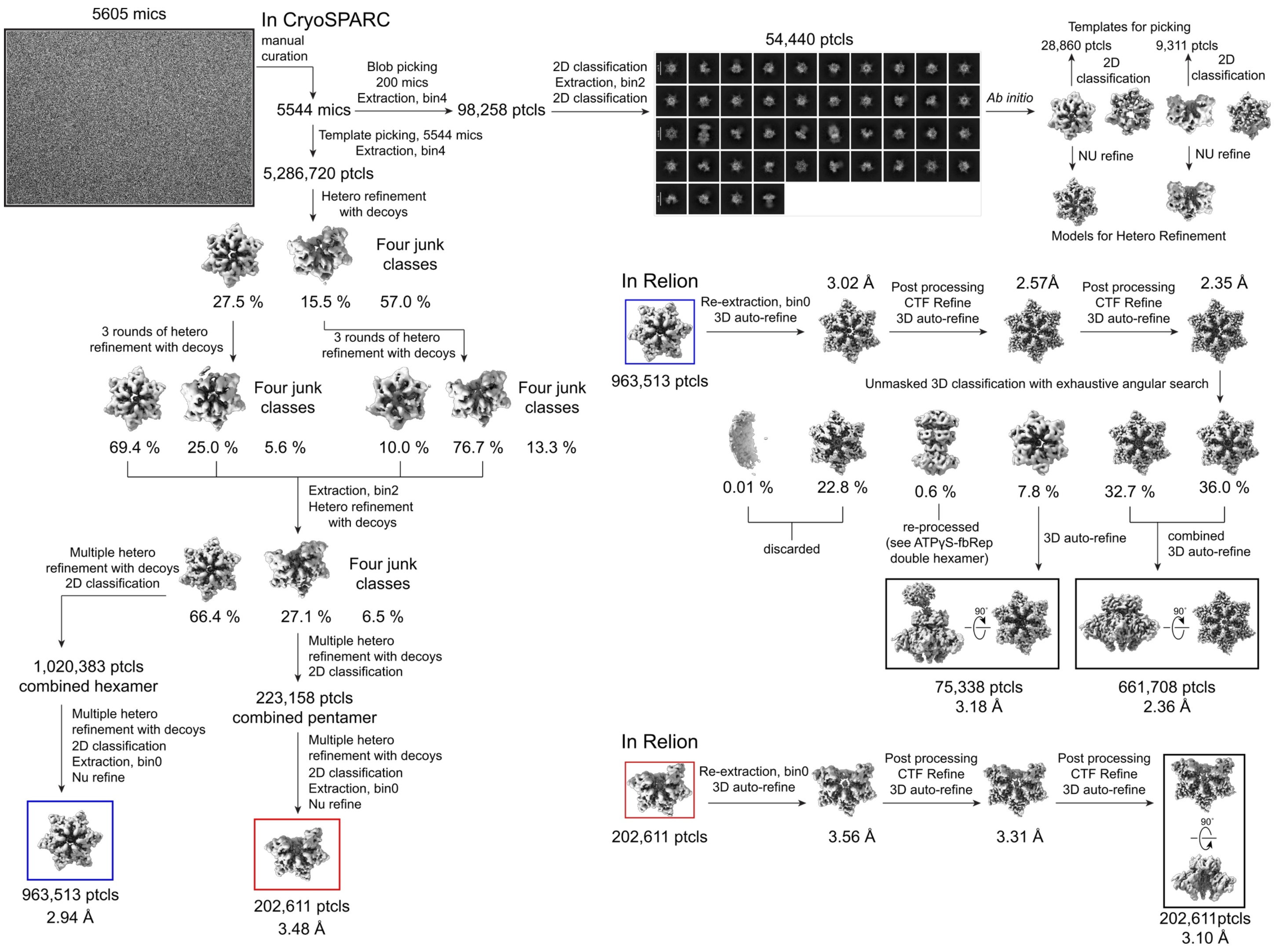


**Supplemental Figure 4.** Cryo-EM data processing workflow of the fbRep•ATPγS dataset. Schematic diagram of data collection, initial data processing, classifications, and refinements used to obtain the final maps (see Methods for details). The orthogonal views of the final maps are shown in black boxes. Resolutions associated with the final maps were calculated in CryoSPARC at FSC threshold of 0.143.

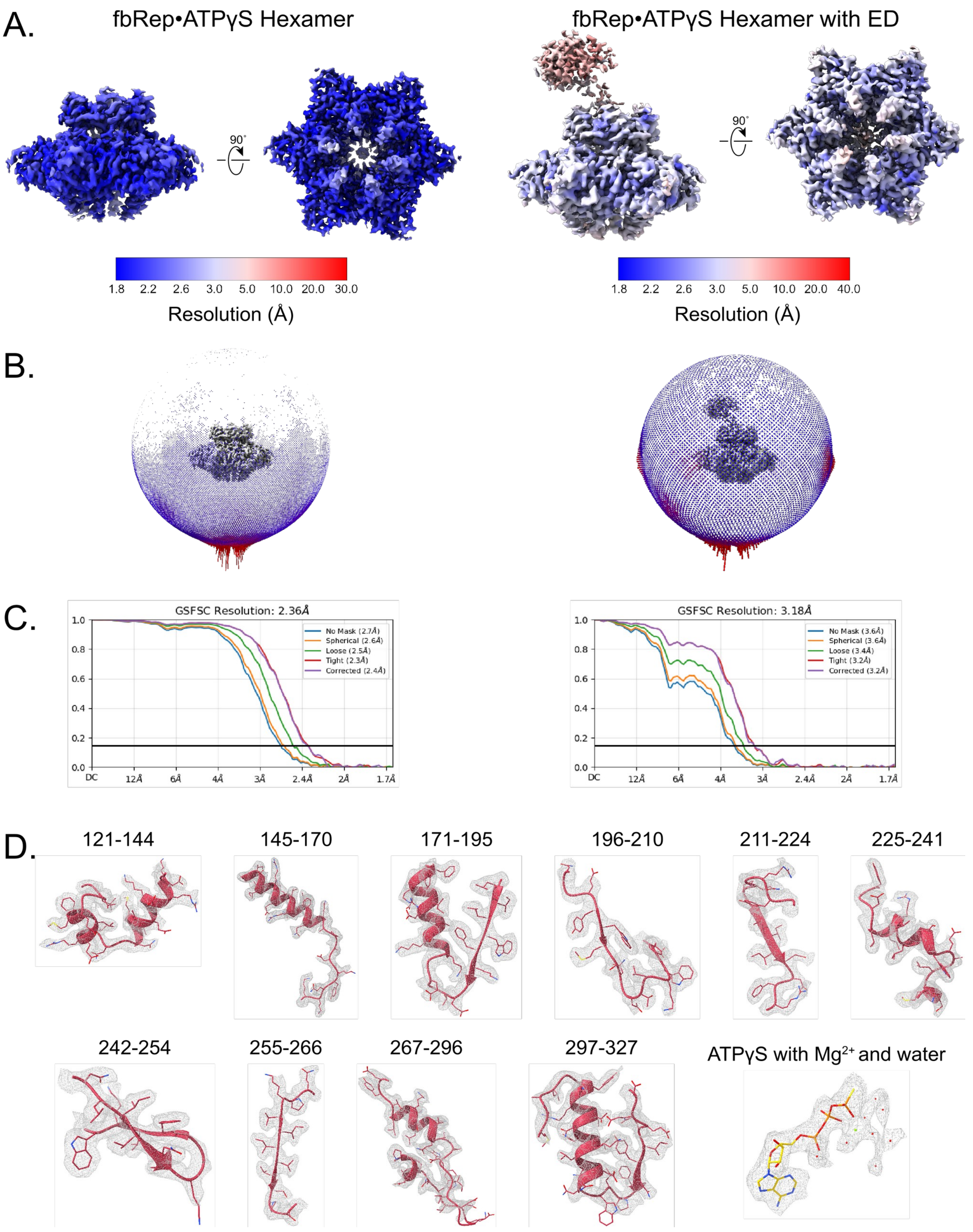


**Supplemental Figure 5.** Analysis of cryo-EM data quality of fbRep•ATPγS hexamers. A. Orthogonal views of 2.36 Å fbRep•ATPγS hexamer and 3.18 Å fbRep•ATPγS hexamer with ED, colored based on estimated local resolution as calculated in CryoSPARC. B. Angular distribution of particles in the final refinement of fbRep•ATPγS hexamer maps. C. Map FSC curves at 0.143 threshold. D. Representative sidechain densities of 2.36 Å fbRep•ATPγS chain A and ATPγS binding site contoured at 4.6σ. Densities for each chain and ATPγS binding site are nearly identical.

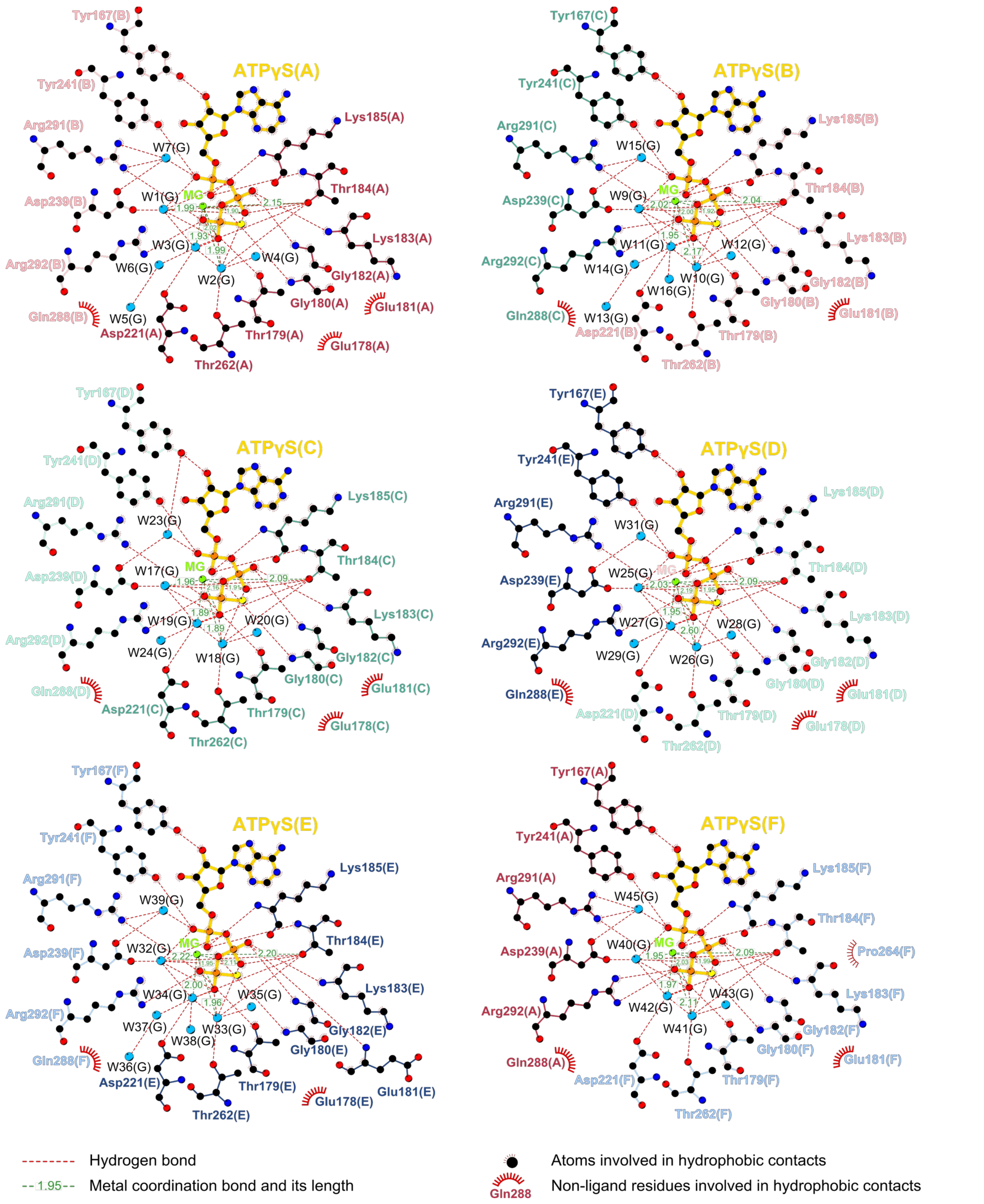


**Supplemental Figure 6.** Ligplot+ diagrams of hydrogen bonding and hydrophobic interactions of ATPγS. Hydrogen bonding and hydrophobic interactions of ATPγS in 2.36 Å fbRep•ATPγS hexamer model were calculated in Ligplot+ v2.2 using HBPlus and default parameters. ATPγS and protein side chains are shown in ball-and-stick representation, with ATPγS colored in yellow. Hydrogen bonds are shown as red dashed line while hydrophobic contacts are represented by red spoked arcs. Magnesium ion coordination bonds and its length, calculated in UCSF ChimeraX based on center-center distance parameter value of ≤ 3.10, are also shown and represented as green dashed lines.

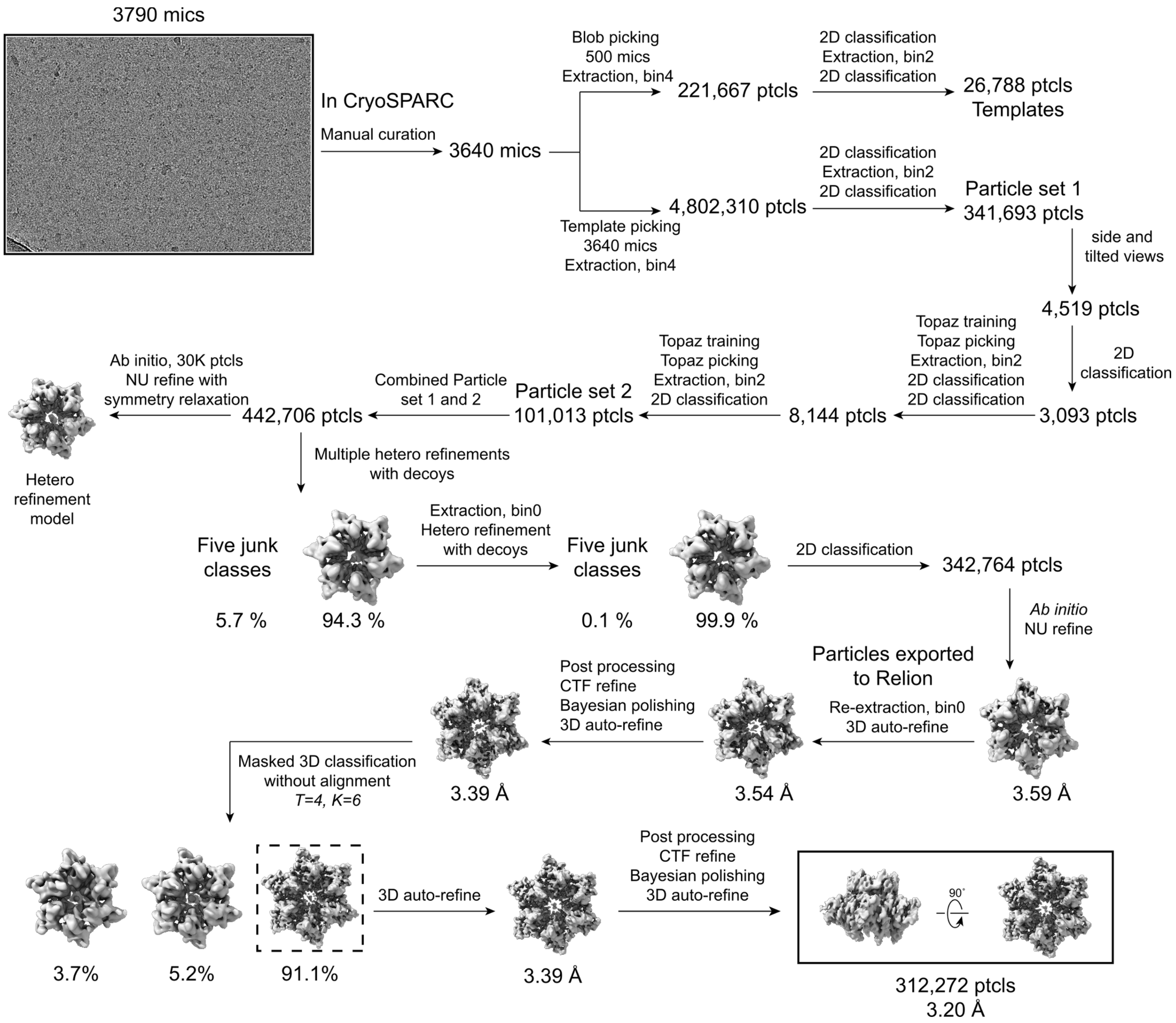


**Supplemental Figure 7.** Cryo-EM data processing workflow of the fbRep•ADP dataset. Schematic diagram of data collection, initial data processing, classifications, and refinements used to obtain the final map (see Methods for details). Resolution associated with the final map was calculated in CryoSPARC at FSC threshold of 0.143.

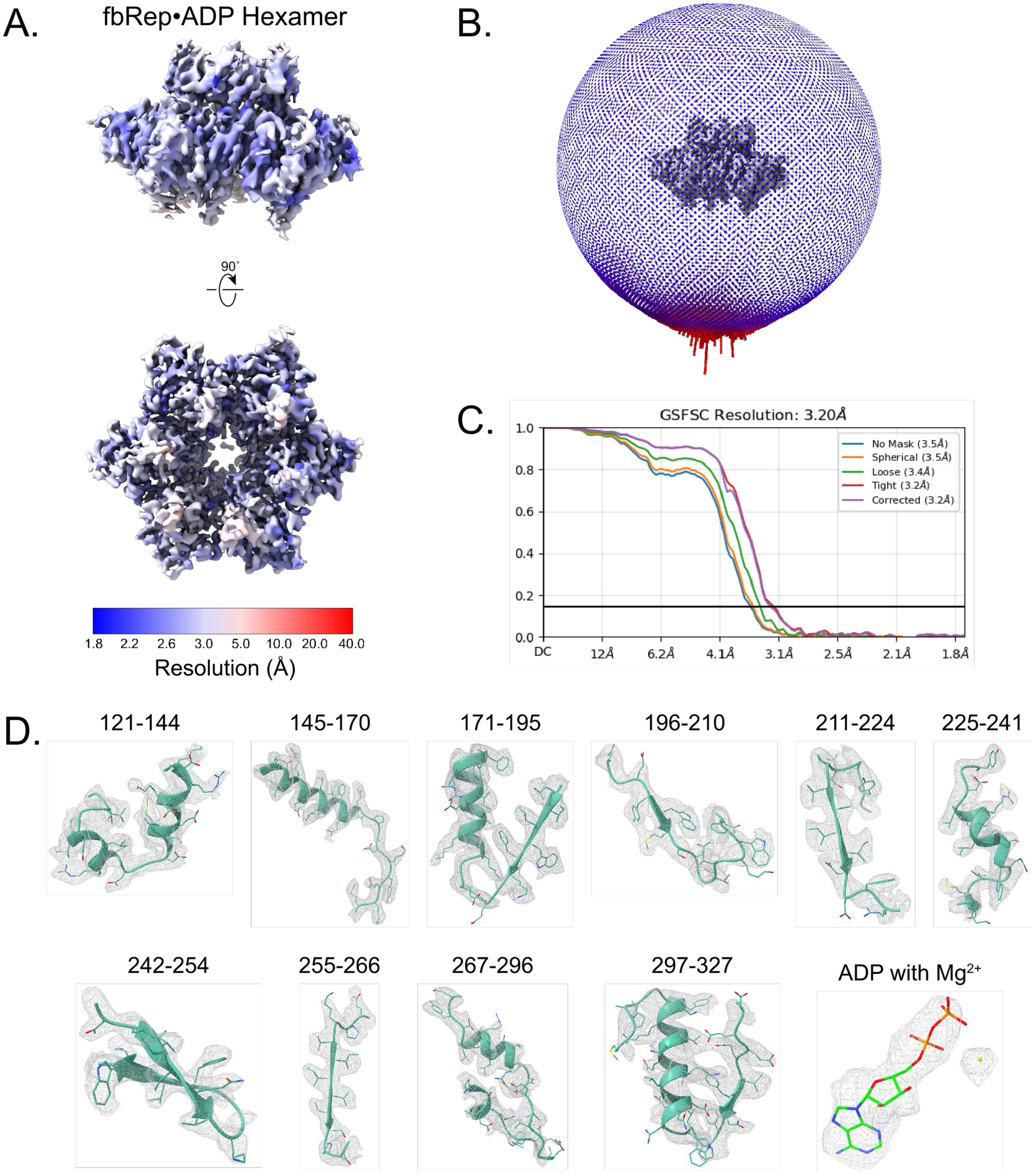


**Supplemental Figure 8.** Analysis of cryo-EM data quality of fbRep•ADP hexamer. A. Orthogonal views of 3.20 Å fbRep•ADP hexamer map, colored based on estimated local resolution as calculated in CryoSPARC. B. Angular distribution of particles in the final refinement of fbRep•ADP hexamer map. C. Map FSC curve at 0.143 threshold. D. Representative sidechain densities of 3.20 Å fbRep•ADP chain C and ADP binding site contoured at 5.8σ. Densities for each chain and ADP have subtle differences.

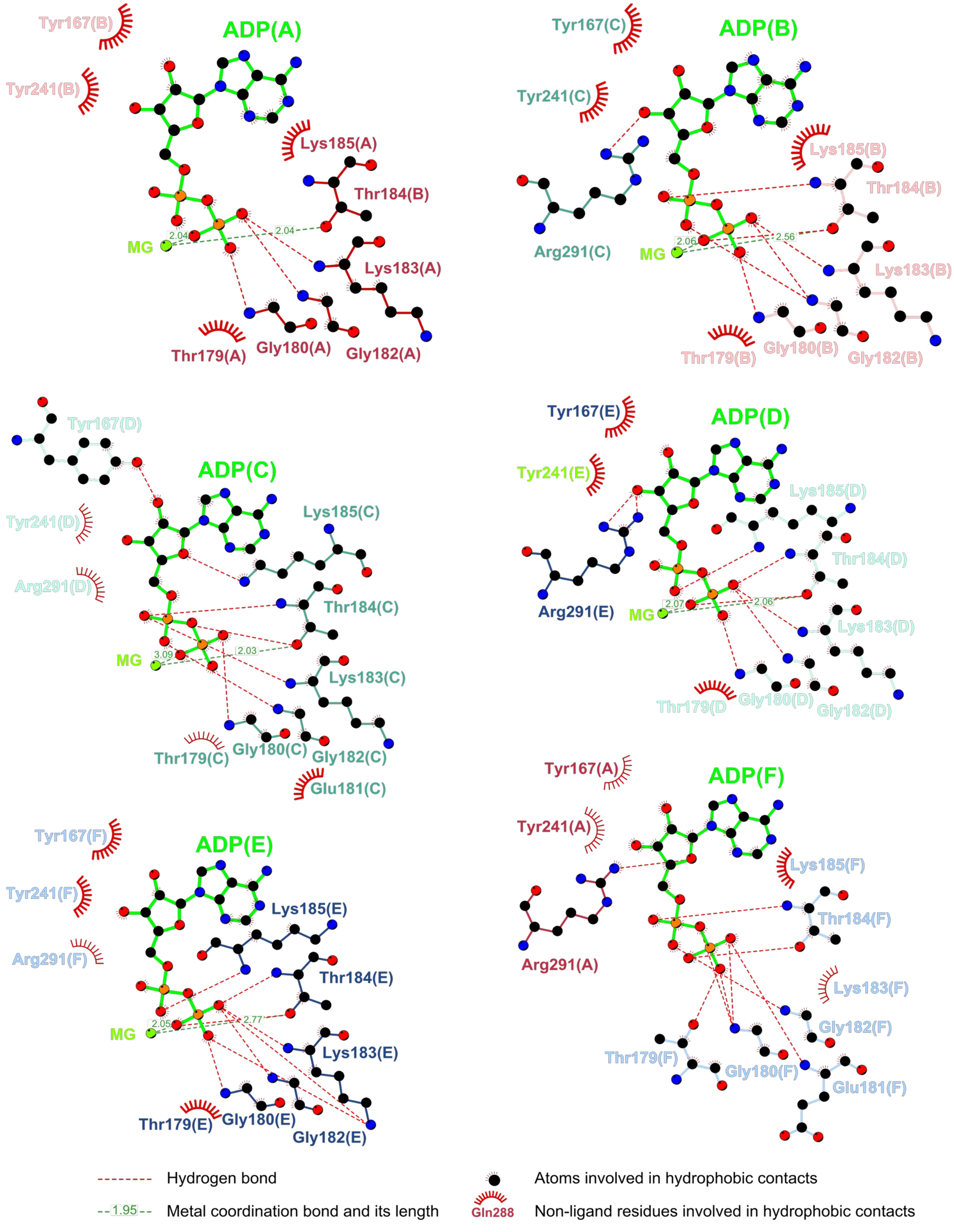


**Supplemental Figure 9.** Ligplot+ diagrams of hydrogen bonding and hydrophobic interactions of ADP. Hydrogen bonding and hydrophobic interactions of ADP in 3.20 Å fbRep•ADP hexamer model were calculated in Ligplot+ v2.2 using HBPlus and default parameters. ADP and protein side chains are shown in ball-and-stick representation, with ADP colored in lime green. Hydrogen bonds are shown as red dashed line while hydrophobic contacts are represented by red spoked arcs. Magnesium ion coordination bonds and its length, calculated in UCSF ChimeraX based on center-center distance parameter value of ≤ 3.10, are also shown and represented as green dashed lines.

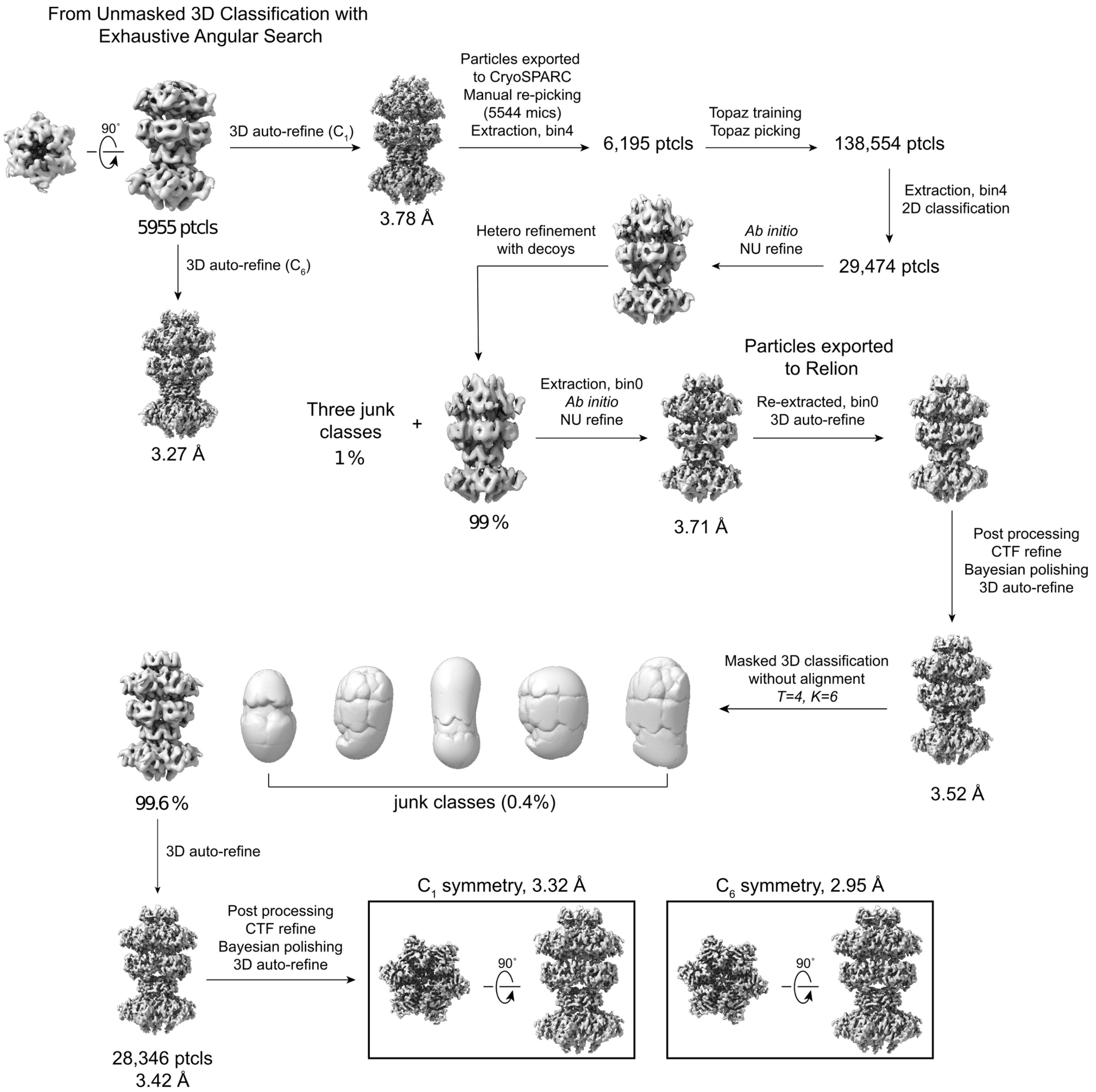


**Supplemental Figure 10.** Cryo-EM data processing workflow of the fbRep•ATPγS double hexamer. Schematic diagram of data reprocessing, classifications, and refinements used to obtain the $C_1$ and $C_6$-symmetrized final maps (see Methods for details). The orthogonal views of the $C_1$ and $C_6$-symmetrized final maps are shown. Resolutions associated with the final maps were calculated in CryoSPARC at FSC threshold of 0.143.

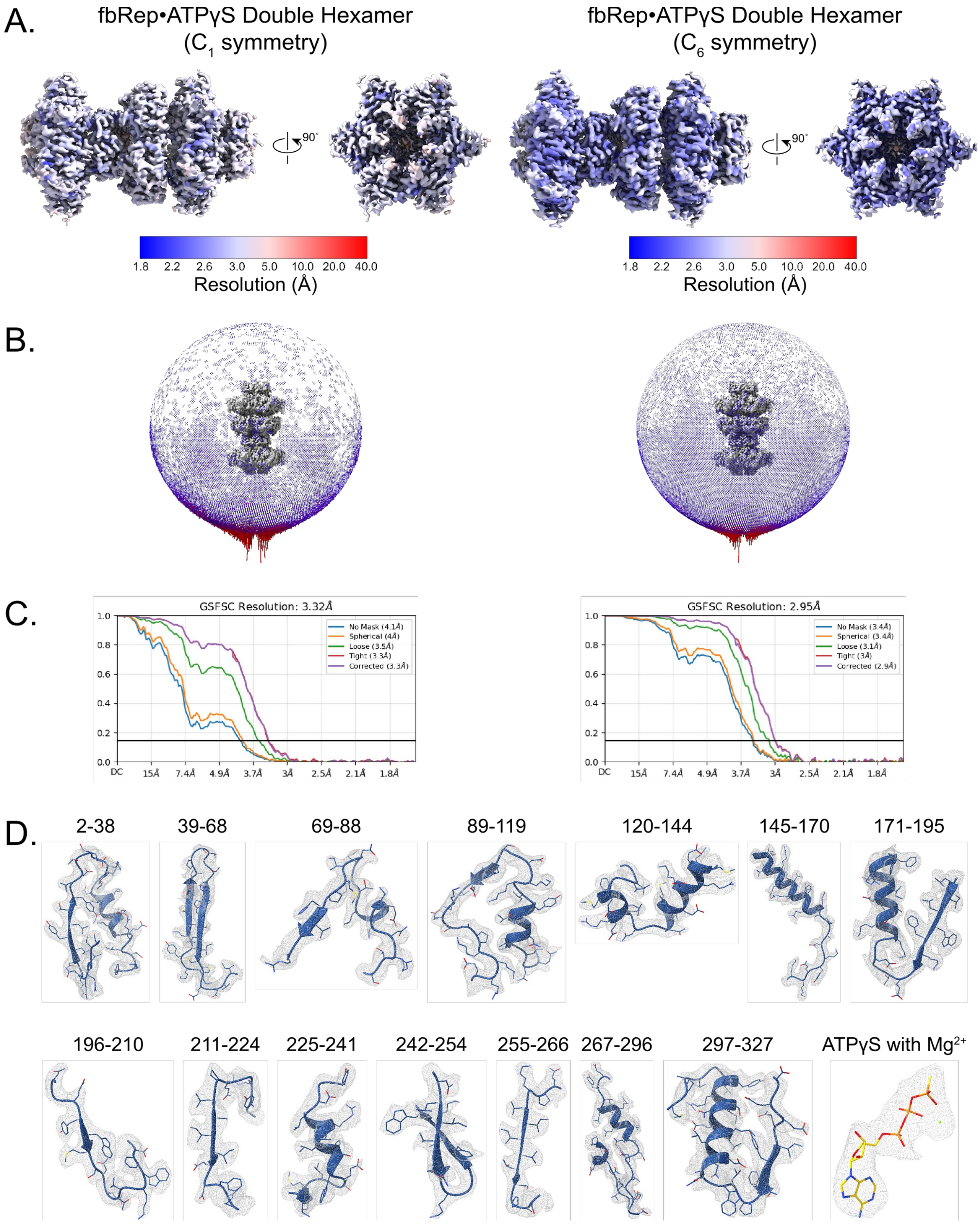


**Supplemental Figure 11.** Analysis of cryo-EM data quality of fbRep•ATPγS double hexamer. A. Orthogonal views of 3.32 Å fbRep•ATPγS double hexamer ($C_1$ symmetry) and 2.95 Å fbRep•ATPγS double hexamer ($C_6$ symmetry), colored based on estimated local resolution as calculated in CryoSPARC. B. Angular distribution of particles in the final refinement of fbRep•ATPγS double hexamer maps. C. Map FSC curves at 0.143 threshold. D. Representative sidechain densities of 3.32 Å fbRep•ATPγS double hexamer chain E and ATPγS binding site contoured at 5.4σ. Densities for each chain and ATPγS binding site are nearly identical.

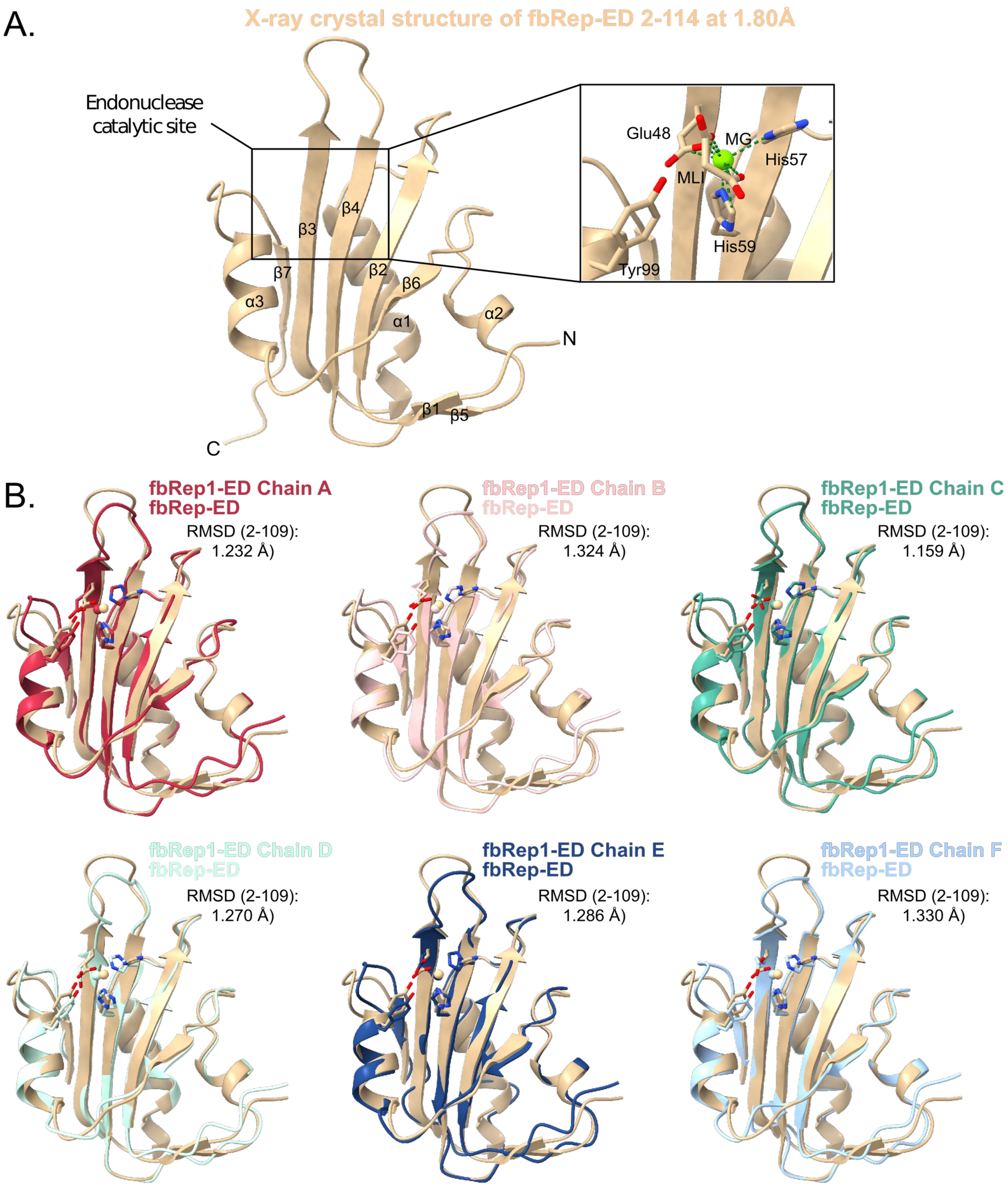


**Supplemental Figure 12.** Structure of fbRep endonuclease domain. A. X-ray crystal structure of fbRep endonuclease domain 2-114 (fbRep-ED) with $Mg^{2+}$ ion resolved at 1.80 Å resolution. The endonuclease domain contains a central five-stranded anti-parallel β-sheet surrounded by α-helices and disordered loops. The α3 helix contains the catalytic tyrosine at position 99 with its side chain pointing towards the metal coordination site. Magnesium is coordinated by Glu48 (β3), His57 (β4), His59 (β4) and a malonate ion (MLI). The coordination of $Mg^{2+}$ by malonate ion likely represents a key interaction of ssDNA phosphate backbone during catalysis. B. Superposition of X-ray crystal structure of fbRep-ED and cryo-EM structure of fbRep1 EDs. Structural alignments of the EDs (2-109) were done in UCSF ChimeraX with matchmaker tool using default parameters. The overall Cα r.m.s.d of 108 paired atoms are indicated.

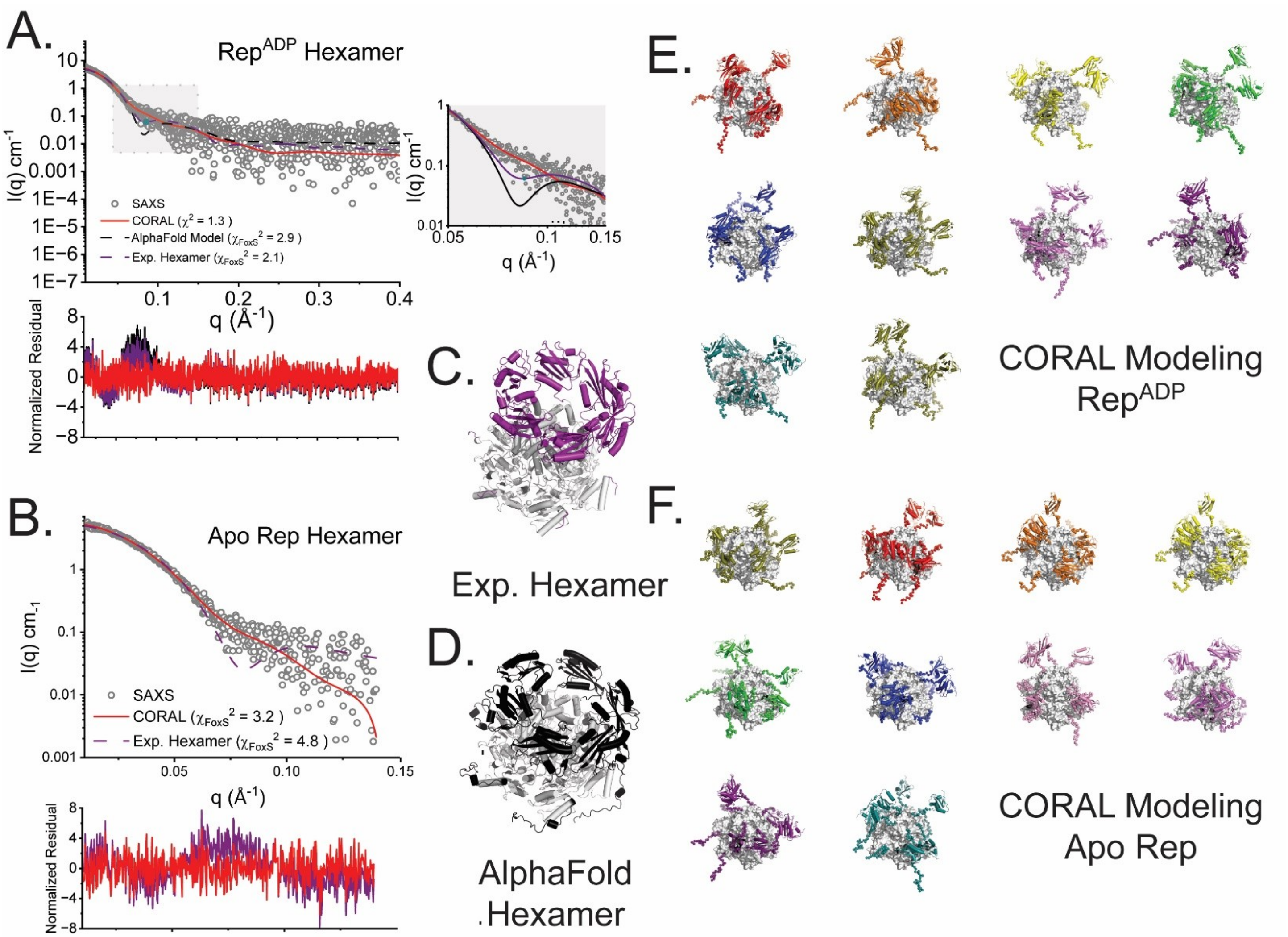


**Supplemental Figure 13. CORAL Modeling.** CORAL analysis (5) of Rep hexamers in the presence and absence of ADP. CORAL employs a rigid body approach to optimize atomistic models against experimental SAXS data. In this approach, missing and flexible atomic inventory are represented as beads in coarse grain fashion and flexibly fit. Shown in **A** are the experimental SAXS data for RepADP, and in **B** the experimental SAXS data for apo Rep (see **Table 1**). In both plots, gray circles represent the experimental data in a log–log plot, where intensity I is plotted as a function of q. In this analysis, two hexamer models containing the N-terminal endonuclease domains (ED) were initially considered: the experimental hexamer determined in these studies (ED domains colored purple, **C**), and an Alphafold3 (AF3, (6, 7)) calculated model (ED domains colored black, **D**). In the ADP state (**A**), the experimental hexamer (purple dotted lines) showed discrepancies with the experimental data between $0.05 < q < 0.1$ ($\chi_{FoxS}^2$ = 2.1 vs ADP, 2.9 vs APO), and similar discrepancy was apparent with the AF3 derived model (black dotted line, $\chi_{FoxS}^2$ = 2.9). When missing atomic inventory (residue 323-354) and the N-terminal domains were modelled using the CORAL approach, this discrepancy is relieved (representative fits shown in red, $\chi_{FoxS}^2$ = 1.31-1.34 for ADP-bound (n=10) and $\chi_{FoxS}^2$ = 3.21-3.32 for apo (n=10)) (8). In all cases, random positioning of the EDs in proximal and distal configurations is observed. A gallery of representative calculations (n=10) is shown in **E** for the ADP-bound form of hexameric Rep and **F** for apo Rep. Atomic inventory not represented in available crystallographic models is shown as beads and fit flexibly. Residues 120-323 (white) were fixed in these calculations. Figures were rendered using PYMOL.
